# Proximity to Explosive Synchronization Determines Network Collapse and Recovery Trajectories in Neural and Economic Crises

**DOI:** 10.1101/2024.11.28.625924

**Authors:** UnCheol Lee, Hyoungkyu Kim, Minkyung Kim, Gabjin Oh, Ayoung Park, Pangyu Joo, Dinesh Pal, Irene Tracey, Catherine E. Warnaby, Jamie Sleigh, George A. Mashour

**Author notes:** The authors contribute to the study equally as the first author. Research Institute of Slowave Inc., Seoul; Center for Neuroscience Imaging Research, Institute for Basic Science (IBS), Suwon, Republic of Korea.

## Abstract

When complex systems move away from criticality—a balance between order and chaos—they are no longer optimized. Furthermore, when criticality is lost too quickly, or recovery is delayed, system damage can result. However, the mechanism for these abnormally fast or slow critical transitions remains unknown. Here, we show that the proximity of a complex network to explosive synchronization (ES), a first-order phase transition, determines the trajectories of criticality loss and recovery after perturbations. Our computational models revealed characteristic dynamics based on network proximity to ES, enabling us to infer network phase transition types from empirical data and predict criticality transition patterns. We validated our predictions using empirical data from the human brain under anesthesia and the stock market during an economic crisis, demonstrating that early and prolonged recoveries can be systematically predicted. This study has implications for designing resilient networks that withstand perturbations and recover quickly.

## Main Text

Rapid collapse during a crisis or delayed recovery afterwards can cause severe damage to systems. For example, excessive sensitivity to or prolonged recovery from anesthesia—a significant perturbation to brain networks— can lead to patient complications^1^. Similarly, a rapid market crash or prolonged economic recovery can have lasting adverse effects on a country’s economy^2^. Understanding the mechanisms of early and delayed transitions in complex dynamic systems, such as the brain and financial markets, is thus crucial for measuring, predicting, and correcting abnormal transitions.

We first define normal states and crises to study system collapse and recovery during crises. Many complex systems in nature operate in a critical state, at the edge of phase transitions, exhibiting typical properties such as scale invariance, long-range correlation, large autocorrelation, and high susceptibility to external perturbations^3^. These properties underpin system capacities for energy efficiency, information integration, spatiotemporal memory, and flexible adaptation. Thus, a critical state is considered the “sweet spot” for emerging high-order functions and complexity in systems^4-7^. However, deviation from a critical state disrupts these abilities. In this study, we define a deviation from criticality as a system crisis and consider the times taken to lose and restore a critical state as surrogates for the times to system collapse and recovery during crises.

When external perturbations are applied, the time to lose and regain criticality is largely determined by the stability of each system’s critical state, the system’s self-organization processes to maintain the critical state, and, most fundamentally, the type of phase transition^8-10^. Phase transitions describe the process of changing from one state to another, with the critical point being the specific condition at which this change occurs. The nature of the change at the critical point can be simply categorized as either first-order or second-order^11^. For example, the transition from water to ice is a first-order phase transition marked by abrupt change, while the transition from a ferromagnetic to a paramagnetic state is a second-order phase transition characterized by gradual change. Therefore, if we could infer a system’s phase transition type, we could anticipate whether it will undergo abrupt or gradual change near critical points. However, there is a paucity of methods to make such inferences, especially from empirical data, which hinders our ability to predict abnormal transitions during crises in real-world systems.

Explosive synchronization (ES) is a first-order phase transition in complex networks characterized by a sudden transition from incoherence to complete synchronization. Recent studies argue that parameters like heterogeneous frequency and degree distribution, higher-order connectivity, or adaptive feedback can generally induce ES^12-14^. Furthermore, simply modulating a few nodes in the complex network can change the network’s phase transition type from non-ES (second-order and gradual) to ES (first-order and abrupt), or vice versa^15-17^. These network connectivity and feedback parameters have parallels in the variation in the strength of inter- and intra-regional brain functional connectivity patterns and in financial networks of assets and investments. Despite fundamental differences in details—the brain is a biological network of neurons and other cells mediating cognition and action, while financial markets are networks of institutions managing capital flow—both systems are thought to operate near critical points in non-equilibrium states. Therefore, we predict that the universality of ES and its characteristic transition patterns near critical points could apply to both systems during crises.

Moreover, as illustrated in Figure 1 A, every network can be positioned along a spectrum ranging from close to distant ES proximity in parameter space. Networks with closer ES proximity may exhibit behavior more akin to first-order phase transitions near critical points (Figure 1 B). Methods based on critical slowing-down phenomena have been widely applied to detect early warning signs of critical transitions in various systems, including ecological, meteorological, and financial systems^18-20^. Similarly, methods based on a universal scaling law have been developed to measure the deviation of complex systems from criticality, thereby differentiating altered brain states from conscious brains^21-24^. However, inferring the system’s proximity to ES when near criticality remains unexplored. This study focuses on inferring a network’s proximity to ES at a critical point, aiming to identify characteristic dynamics of networks with varying ES proximities and investigate the determinant roles these dynamics play in entering or exiting crises.

**Fig 1.**
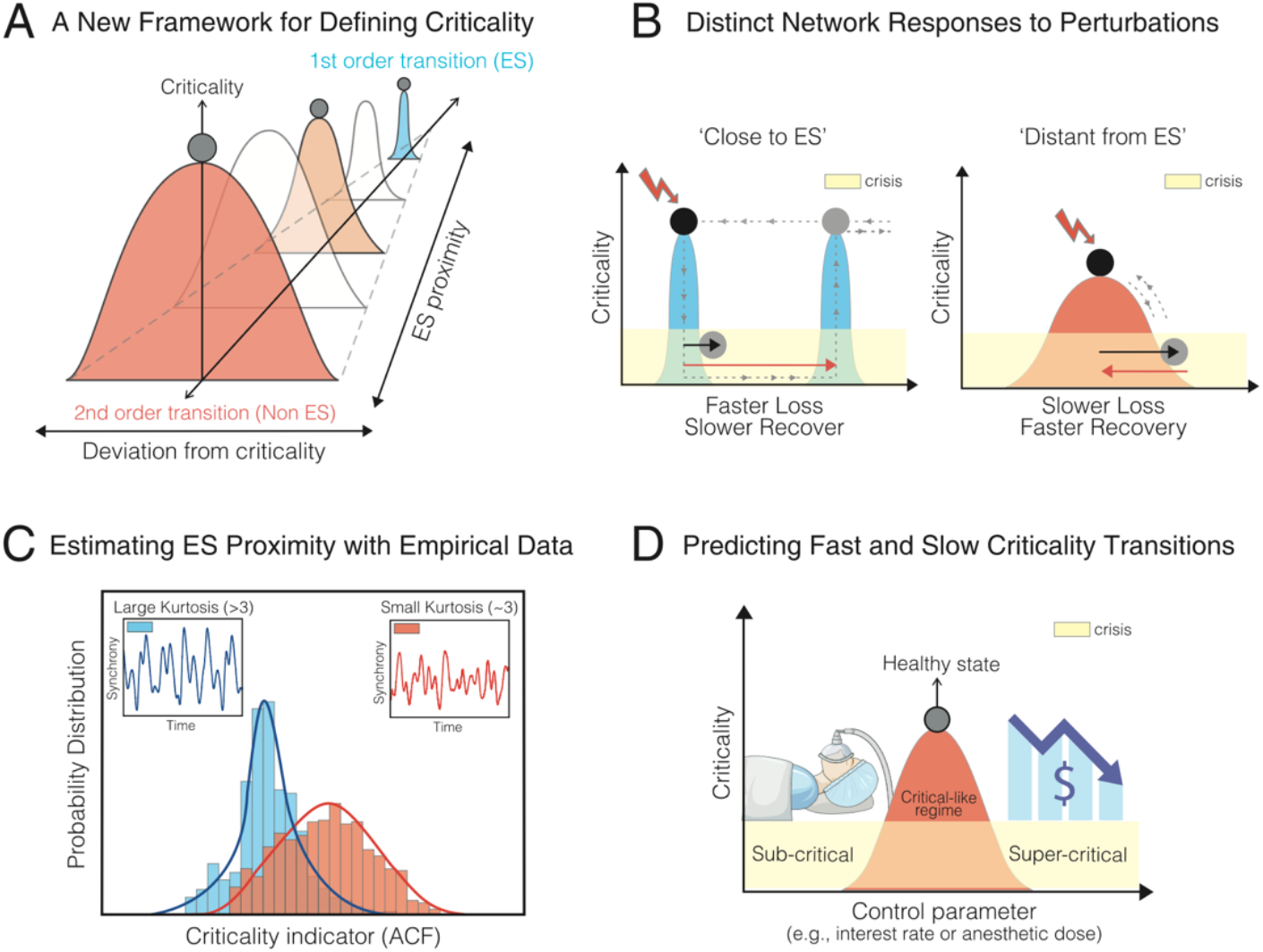
Network explosive synchronization (ES) proximity, resilience, and recovery under perturbation. (A) Networks at healthy and normal states stay near critical points, and the network crisis is characterized by its deviation from the critical point. Individual networks have diverse proximities to ES, i.e., first-order phase transition, resulting in larger interindividual variability in criticality loss and recovery. (B) Response behaviors to perturbations differ according to ES proximity: networks with ‘close’ ES proximity are more susceptible at critical points and show greater internal resistance, manifested as hysteresis, leading to faster criticality loss and slower recovery. Black and red arrows mark approximate times of criticality loss and recovery, respectively. (C) Networks with ‘close’ or ‘distant’ ES proximities exhibit distinct dynamics at critical points or critical-like regimes, which can be estimated through time series signals. (D) Real-world complex networks, such as the human brain and financial networks, operate near critical points in healthy and normal states. When these systems deviate from criticality, as during anesthesia-induced neuronal crises or financial crises like 2008, pre-crisis ES proximity significantly influences the rates of consciousness and market collapse and recovery.

This study investigates (1) Whether networks with distant or close ES proximities have distinctive network dynamics at their critical points, enabling us to infer network proximity to ES based on their time series data. (2) Whether a network’s ES proximity, measured before a crisis, can predict rapid or prolonged network collapse and recovery during and after a crisis. (3) How network structure and dynamics influence ES proximity and transition patterns under external perturbation. (4) Whether these methods can be applied to real-world complex networks, such as brain and financial networks during neural and economic crises, to predict rapid or prolonged transitions. Figure 1 illustrates the schematic overview of this study.

### The proximity of a complex network to explosive synchronization determines rapid or prolonged transition near a critical point: a computational model study

We examined how a network’s proximity to ES affects dynamics near critical points and the response to perturbations. The conventional Stuart-Landau model, a simplified representation of an oscillator near a Hopf bifurcation, has been used to study network dynamics near and far from critical points^25^. We employed a modified Stuart-Landau model with two key parameters: one varying ES proximity^26^ and the other varying perturbation strength. This model allows us to characterize network dynamics at critical points for varying ES proximities and analyze their responses, quantifying the time for critical state loss and recovery. We further explored how different network topologies—random, scale-free, small-world—affect these dynamics.

Our model adjusts ES proximity by manipulating the adaptive feedback strength (Z) among interconnected nodes. The adaptive feedback suppresses synchronization and acts as internal resistance to changes in synchronization. It competes with the force promoting synchronization as the coupling strength among nodes (the other main control parameter) increases. When these two forces are balanced at a critical point, the network becomes highly susceptible to small perturbations, leading to abrupt transitions—this is the core mechanism of ES. We hypothesize that a network with closer ES proximity loses its baseline critical state more easily due to high susceptibility and has prolonged recovery because of significant internal resistance during state transitions, often manifested as hysteresis.

To test this hypothesis, we constructed models comprising 78 interconnected Stuart-Landau oscillators representing a real-world complex network (a diffusion tensor image-informed human brain network), and 1,000 interconnected Stuart-Landau oscillators for small-world, random, and scale-free networks with 4,000 structural links. Each oscillator simulates node activity and interactions with linked nodes. We ran 100 simulations for each of the seven ES proximities, varying initial conditions. To identify each network’s critical point, we tested the pair correlation function (PCF) and autocorrelation function (ACF) of the instantaneous order parameters. The PCF measures the variance of collective phase synchronization fluctuations, while the ACF measures the temporal memory of these fluctuations. Both the PCF and ACF reach maximal values at critical points^27,28^. In this simulation study, we used the PCF peak to identify the critical points because it is non-parametric and was better at finding peaks than the ACF in our model.

In Figure 2A, we illustrate maximal PCFs with yellow circles for Z=0 and Z=2. The network with closer ES proximity (Z=2) exhibits a steeper synchronization transition at the critical point than the one with distant ES proximity (Z=0). We analyzed time series signals across various ES proximities and found that networks with closer ES proximities displayed more variability in sequential ACF values (Figure 2B), reflecting intermittent and bistable transitions near their critical points (Figure S1.A-D). This indicates that these networks tend to produce uncommon ACF values, including extremely high and low ones, leading to fat tails in ACF distributions (Figure 2C). The kurtosis of the ACF distribution, which reflects these fat tails, is correlated with the adaptive feedback strength Z. As Z increases and begins to influence network dynamics near the critical point (Z > 2), the kurtosis of the ACF distribution increases significantly (p < 0.05; Tables S1 and S2 for the statistical tests). From Z > 2, the network dynamics exhibit heavy tails (kurtosis greater than 3). This suggests that the kurtosis of the ACF distribution at a critical point could serve as an indicator of ES proximity. We also obtained similar results with the kurtosis of the PCF (Figure S1.E and F). However, since the kurtosis of the ACF aligns more consistently with the real-world data we tested, our presentation was focused on the ACF results.

**Figure 2.**
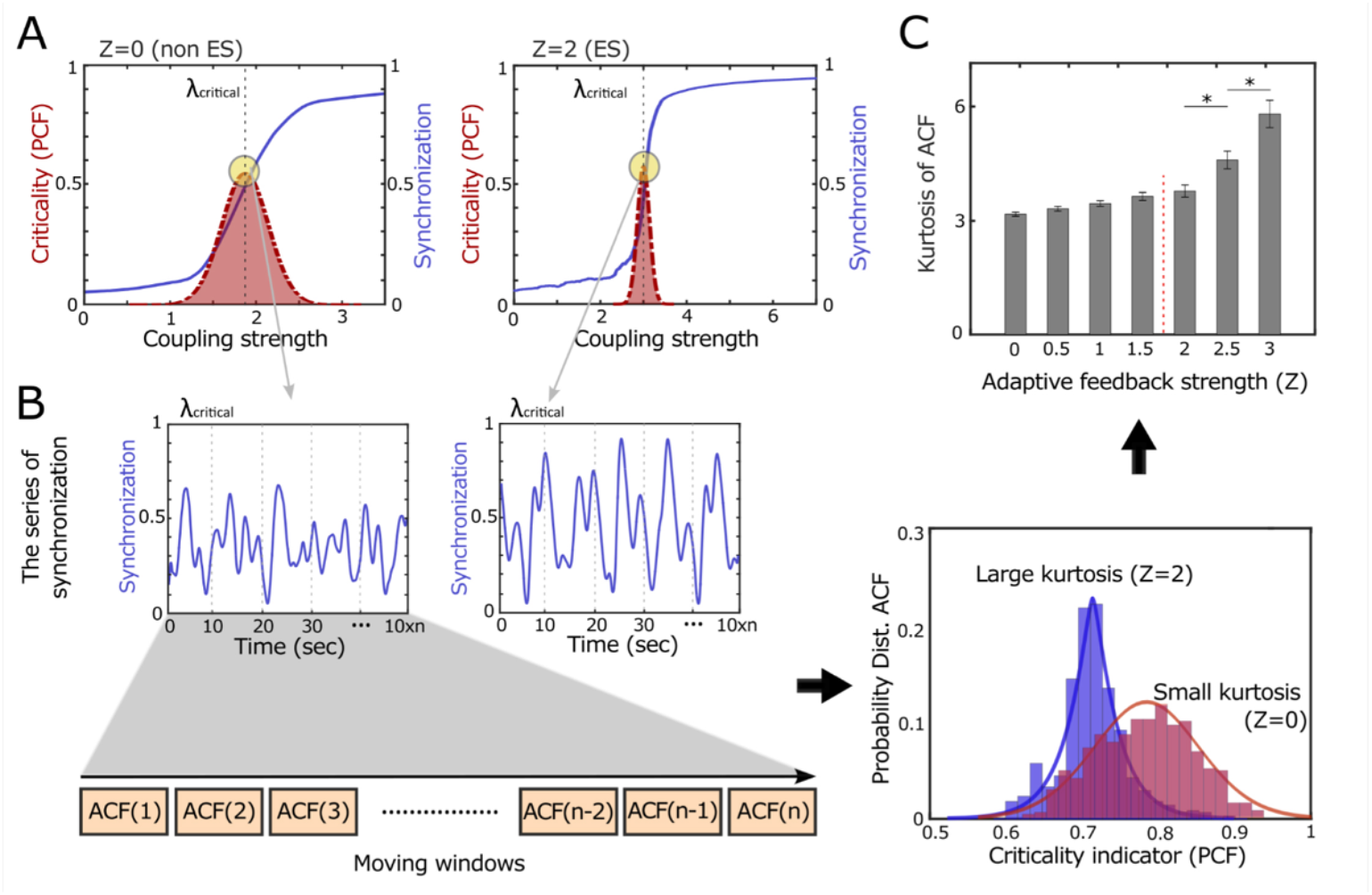
Distinct network dynamics at critical points with varying explosive synchronization (ES) proximity. (A) Networks with close (Z=2) and distant (Z=0) ES proximities exhibit distinctive network dynamics near their critical points. The network with closer ES proximity (Z=2) undergoes a steeper phase transition near its critical point than the network with distant ES proximity (Z=0). Critical points were identified based on the maximal pair correlation function (PCF) of instantaneous order parameters (indicated by circles). The red dashed line indicates the PCF values at each coupling strength. (B) A moving window technique was employed to analyze the variability of the autocorrelation function (ACF) of order parameters, distinguishing the close and distant ES proximities. (C) The ACF distribution of a close ES proximity (Z=2) has a larger kurtosis compared to that of the distant ES proximity network (Z=0). The distinct ACF distributions, which reflect different network dynamics at critical points, demonstrate the potential to estimate the ES proximity using time series data of a complex dynamical network.

To test how networks with different ES proximities respond over time, we introduced external perturbations u(t). Figure 3A illustrates how these perturbations disrupt baseline dynamics at a critical point and how the network returns to its baseline state as the perturbation fades. In our model, the fixed coupling strength at the critical point ensures the network self-organizes and returns to its baseline state after the perturbation ends. We measured the time for the network to deviate from and return to the baseline state by defining a zone as three times the standard deviation of ACF values at baseline. We then calculated correlations between the kurtosis of baseline ACF values before perturbation and the time required for the network to lose and regain its baseline dynamics.

**Figure 3.**
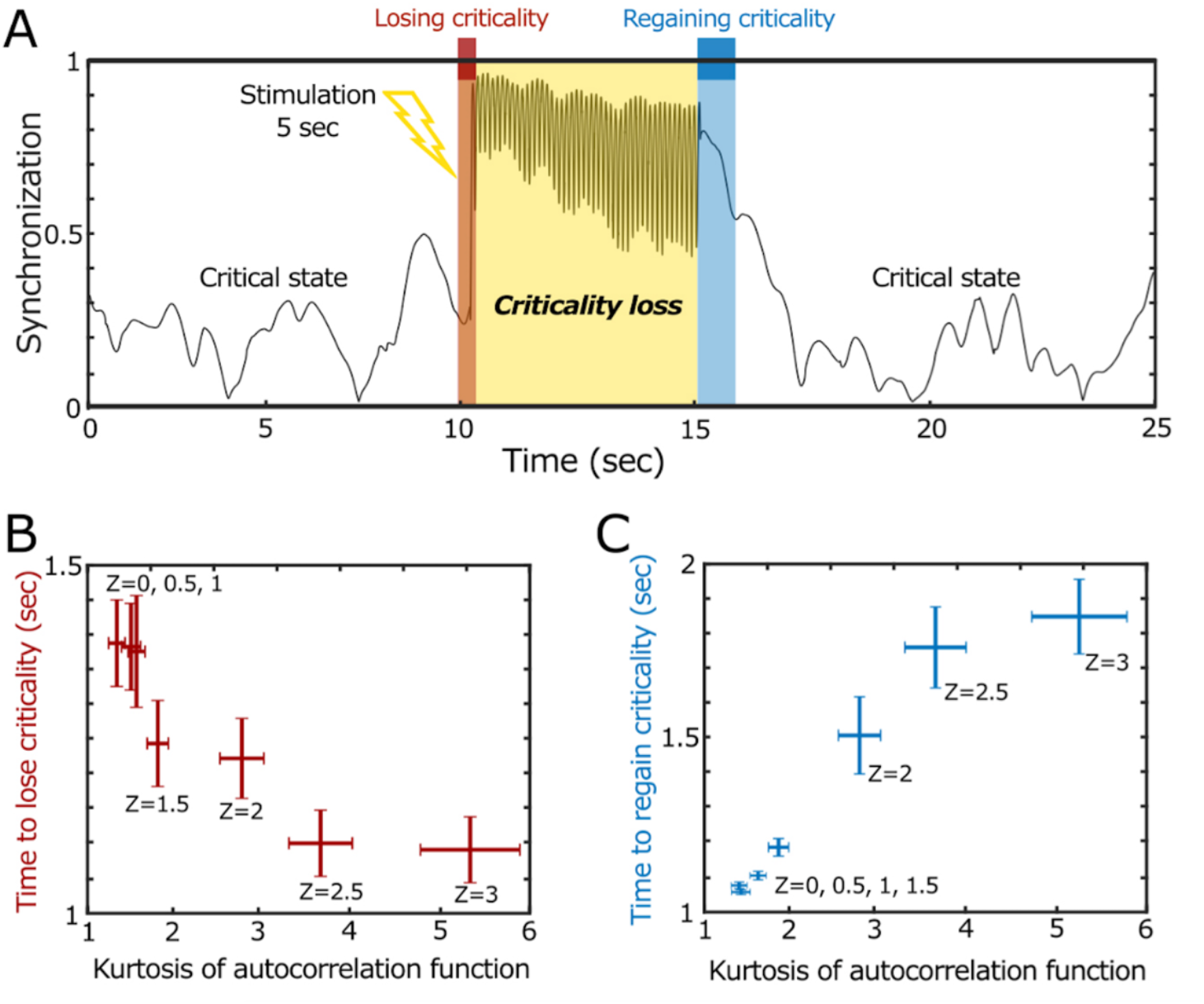
The influence of ES proximity on the critical state loss and recovery under external perturbation. (A) a network with close ES proximity (Z=2) presents characteristic critical state loss and recovery patterns. Under external perturbation, the network loses and regains its baseline critical state. The times to lose (red zone) and regain (blue zone) the baseline states were determined by the times when the network’s ACF values cross over and return to within three standard deviations of the ACF values in the baseline state. (B) The kurtoses of baseline ACF distributions are negatively correlated with the times to baseline critical state loss, indicating that networks with closer ES proximity (a larger Z) are more prone to faster critical state loss. (C) Conversely, the kurtoses of baseline ACF distribution are positively correlated with the recovery time. A network with closer ES proximity (a larger Z) exhibits slower critical state recovery. Error bars indicate standard errors of 100 simulations.

Figure 3A shows the response of closer ES proximity (Z=2) to an external perturbation. After a 5-second perturbation at u(t) = 10, the network exhibits an abrupt deviation from baseline followed by gradual recovery. Figures 3B and 3C demonstrate significant correlations between the kurtosis of ACF values and the times taken for critical state loss and recovery. We observed that the kurtosis of baseline ACF is negatively correlated with the time of critical state loss and positively correlated with recovery time. The robustness of the results was confirmed by testing various strengths u(t) (20, 40, 60, 80, and 100, Figure S2) and network structures (random, scale-free, small-world, Figure S3) (Figure S3). This study suggests that networks with closer ES proximity take less time to lose their baseline critical state but require more time to recover after perturbation.

### Proximity to explosive synchronization of brain networks during baseline consciousness determines the temporal course of anesthetic-induced state transitions

General anesthesia represents a neural “crisis” where functional brain networks are perturbed, and the brain deviates from a critical state^29-31^. We hypothesized that early or delayed transitions in loss and recovery of consciousness might be governed not only by the pharmacokinetics of anesthetics but also by the type of phase transition in functional brain networks. According to our model, we expected that the kurtosis of ACF of the electroencephalogram (EEG) recorded in a resting state, which is a surrogate for the ES proximity of the brain network, may determine the temporal course of state transitions at the boundaries of consciousness. Due to the high susceptibility and hysteresis of an ES network, we also expected that brain networks with a large kurtosis of ACF may exhibit a fast loss and slow recovery of consciousness during anesthetic transitions.

To test the hypothesis, we analyzed data from 16 healthy human subjects who had 32-channel EEG recorded during anesthetic state transitions. Figure 4A illustrates the EEG electrodes and signals. Figure 4B presents a single subject’s EEG spectrogram, and the states studied: resting state with eyes closed (10 min), induction period (start of anesthetic delivery to loss of consciousness), unconscious period (loss of consciousness to maximal anesthetic concentration), and recovery period (maximal anesthetic concentration to the recovery of consciousness). Times to loss and recovery of consciousness were determined by behavioral response to verbal command. Despite maintaining the same anesthetic effect-site concentration across subjects, the times of consciousness loss and recovery were largely variable (See y-axes in Figure 4C and D).

**Figure 4.**
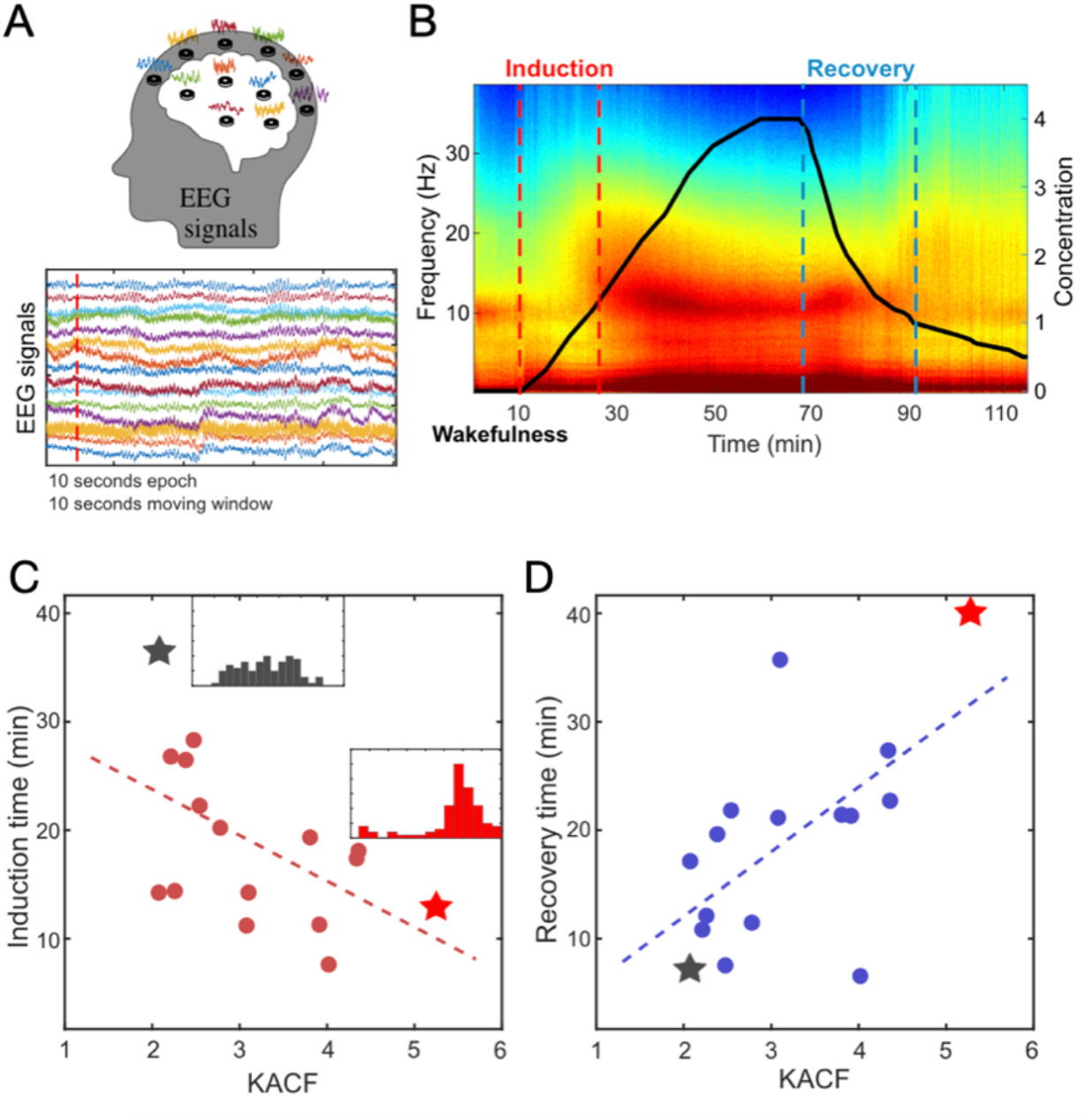
The ES proximity of human EEG in a conscious resting state significantly correlates with the induction time and the recovery time in general anesthesia. (A) Thirty-two channel EEGs during baseline consciousness, induction of anesthesia, unconsciousness, and recovery states were analyzed to test the relationship between ES proximity in conscious brains and fast/slow state transitions induced by the general anesthetic propofol. (B) The spectrogram shows a significant change in the spectral content of the EEG along with state transitions in anesthesia. The anesthetic induction and recovery times were defined by the time intervals between the injection of propofol and the loss of responsiveness (red dotted lines) to a verbal command (induction) and between the end of injection and the recovery of response (blue dotted lines) to a verbal command (recovery). The solid line indicates the modeled effect-site concentration of propofol in the volunteer’s brain. The kurtosis of ACF calculated with the baseline EEG shows a significant negative correlation with the induction times (C) for 16 subjects. Conversely, the kurtosis of the ACF of the baseline EEG positively correlates with the recovery time (D).

We calculated the kurtoses of ACFs using 3-minute clean EEGs from resting states after pre-processing and noise treatments (See Supplementary Method S2 for details). First, we applied band-pass filtering to isolate the alpha-frequency band (8-13Hz) and extracted the instantaneous phases with the Hilbert transform. Using these phases, we calculated the instantaneous order parameters for the EEG signals. To analyze the kurtosis of ACF, we employed a moving window on the sequence of instantaneous order parameters. Specifically, ACFs with a time lag of 50 (reflecting the alpha dynamics at ∼10Hz and a sampling frequency of 500Hz) were computed within 10-second windows with a 5-second overlap. Finally, we calculated the kurtosis of the ACFs from all windows. We then assessed the correlation between the kurtosis of ACFs in resting states and times of consciousness loss and recovery. The results showed significant negative and positive correlations between resting-state kurtosis of ACFs and times to consciousness loss (Spearman correlation coefficient: ρ = -0.67, *p*<0.01) and recovery (Spearman correlation coefficient: ρ = 0.59, *p*<0.01), respectively. These findings indicate that an individual brain’s ES proximity in baseline states, measured by kurtosis of ACFs in EEG networks, significantly influences the pattern of conscious state transitions at the boundary between consciousness and unconsciousness.

### Pre-crisis proximity to explosive synchronization of stock market networks determines the time courses of stock market collapse and recovery associated with the 2008 economic crisis

To determine whether ES proximity is a general principle of state transitions during crises across computational, neurobiological, and economic systems, we examined the 2008 economic crisis. This crisis was triggered by the collapse of the U.S. subprime mortgage market, subsequently causing widespread banking system failure and, eventually, worldwide recession. Using the S&P Compustat Global database, we collected daily stock prices from 39 global equity markets from 2006 to 2010. Countries’ GDPs per capita (USD) are shown on the world map (Figure 5A). Details of data collection and processing are found in the Supplementary Methods (See Table S3).

**Fig 5.**
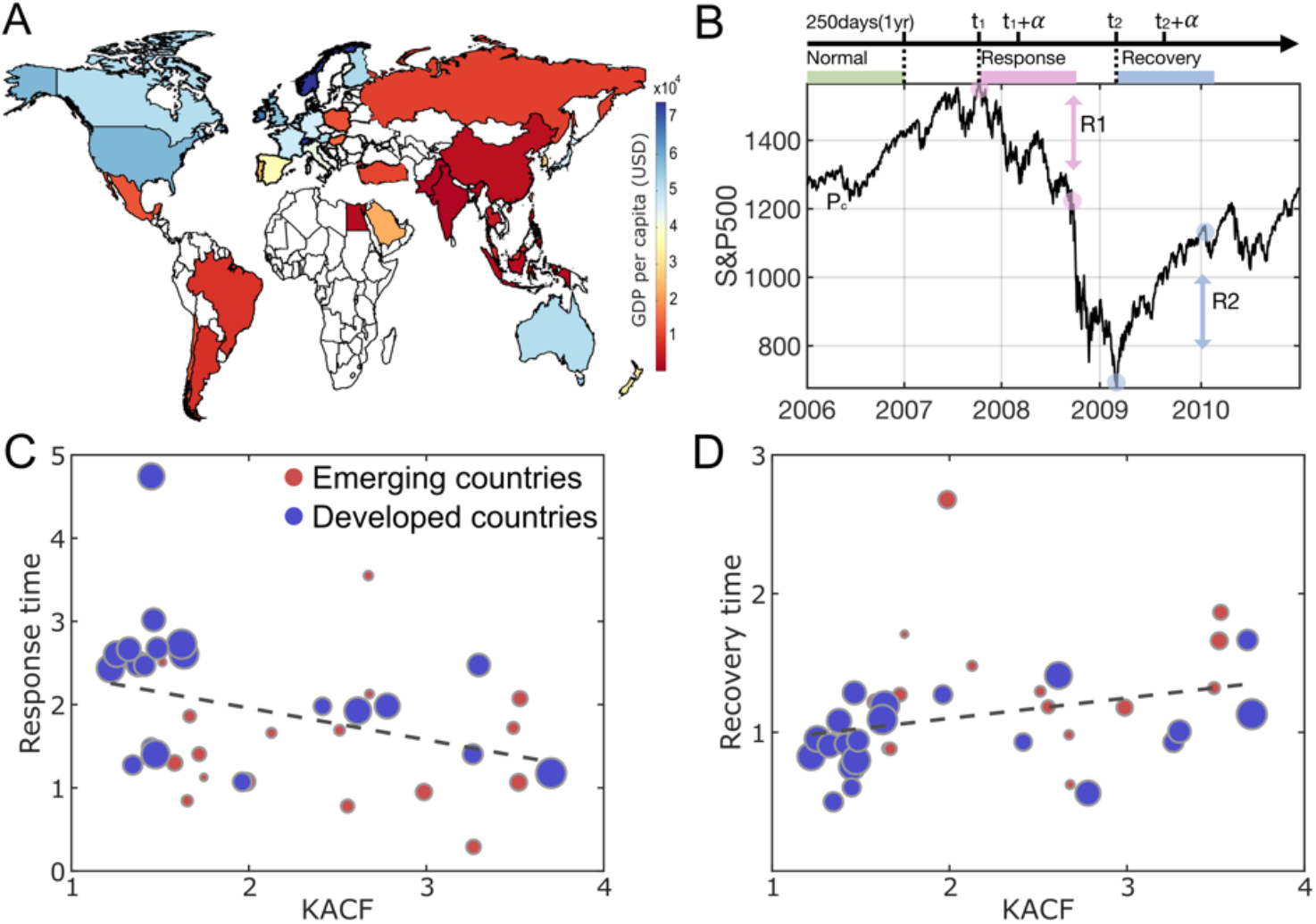
The ES proximities of stock market networks significantly correlate with the rates of market collapse and recovery during the 2008 economic crisis. (A) The thirty-nine countries analyzed in this study are mapped according to their 2006 GDP per capita (USD), with colors ranging from red to blue. (B) The market price of the S&P 500 index underwent a dramatic price change in the 2008-2009 economic crisis, which created natural epochs of baseline (pre-crisis), response (intra-crisis), and recovery (post-crisis) periods. The ES proximity of a stock market network was calculated in the baseline period, and the response and recovery times were calculated with the market collapse (R1) and recovery (R2) rates. (C) and (D) The kurtosis of ACF in the baseline period is negatively correlated with the response time (*ρ* = - 0.40, *p*<0.001), and positively correlated with the recovery time (*ρ* = 0.49, *p*<0.001). Blue and red circles represent developed and emerging countries, respectively, and the marker sizes are scaled by the country’s GDP.

We defined analysis periods for each market: baseline, response, and recovery. Based on studies of the 2007-2009 Subprime Mortgage Crisis^32^, we designated 2006 as the baseline period, preceding the crisis. We tested the robustness of our results by varying the baseline period (See Table S4 for test results). We defined response and recovery periods for each country by calculating the market collapse rate (R1) and recovery rate (R2) during the 2008 crisis, triggered by Lehman Brothers’ collapse in September 2007. Recession periods were identified using recession indicators from the Organization of Economic Cooperation and Development (OECD). We first identified the maximum stock price during the recession period and evaluated the collapse rate over a period α, reflecting the price drop from the maximum. To assess the recovery rate, we identified the minimum price during the crisis and calculated how much the stock price rose from that minimum over the same period α (Figure 5B). Considering the large difference in the price ranges among countries, we normalized market collapse and recovery rates, R1 and R2, for each country (*c*) as follows.

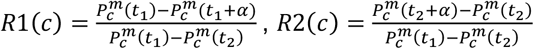

Where 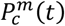 is the stock market index of country *c* and time *t* = 1, 2, 3, … *T*. 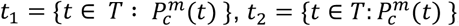, indicate dates of maximal and minimal stock prices, respectively. Because the 39 stock markets have diverse dynamics, we tested several time periods, α = {40, 60, 80, 100, *and*120 *days*}, to calculate collapse and recovery rates. 40 and 120 days correspond approximately to 2 and 6 months, considering market closures. Here, we chose α of 100 days, an appropriate period to reflect the scale of price changes during the crisis. However, the results were not sensitive to α (See Table S5 for test results). Finally, we applied logarithms to the inverse of the market collapse and recovery rates for direct comparison with the EEG study’s induction and recovery time.

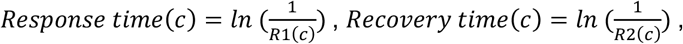

With this transformation, a shorter response/recovery time (i.e., a faster response/recovery rate) corresponds to a shorter induction/recovery time in the EEG study. The financial network analysis showed that the kurtosis of ACFs in the baseline period negatively correlates with the market response times (Spearman coefficient, ρ = - 0.40, *p*<0.001, Figure 5C); conversely, the kurtosis of ACFs positively correlates with market recovery times (Spearman coefficient, ρ = 0.49, *p*<0.001, Figure 5D). In other words, stock markets with higher ES proximity at baseline showed faster market collapse and slower recovery after the crisis. This is consistent with our computational predictions and the empirical neurophysiology of anesthetic state transitions. Moreover, we found the kurtosis of ACF (lag2) significantly correlates with the gross domestic product (GDP) per capita (ρ = -0.36, *p*<0.05) (See Table S6 for the test results). These results indicate that the state transitions in emerging markets, for instance, Thailand, Philippines, and Indonesia, which are classified as Morgan Stanley Capital International (MSCI) emerging markets, are relatively closer to ES than those in developed markets and, therefore, are highly unstable during economic crises.

## Discussion

We investigated the impact of a network’s proximity to explosive synchronization (ES) on the time courses of network collapse and recovery during and after computational, neuronal, and financial crises. Our computational model showed that networks with different ES proximities exhibit characteristic dynamics at their critical points, characterized by the kurtoses of their autocorrelation function (ACF) values. We demonstrated that the kurtosis of ACF values before perturbation is negatively correlated with the time to critical state loss and positively correlated with the time to critical state recovery. This finding suggests that the characteristic transition properties of first-order phase transition determine the time courses during network crises. Specifically, networks with high ES proximity tend to lose their critical state faster and take longer to recover. This relationship was confirmed with diverse model networks (scale-free, small-world, and random) and real-world networks (human brain networks during anesthesia and stock market networks during the 2008 global economic crisis).

### Explosive synchronization proximities of the human brain and stock market networks

Although the brain and financial networks are composed of distinct elements and operate on different principles, both share features as nonequilibrium systems where flows of energy, matter, and information are continuously introduced. The brain is a hierarchical network of cells processing signals for sensory perception, motor control, cognition, and emotions, while financial networks interconnect banks, investors, and institutions through assets and investments to uphold market stability and efficiency. Despite their differences, both networks reside near their critical states under normal conditions^3,4,5,9,33-35^. We hypothesized that each network’s response to significant perturbations depends on its inherent proximity to a first-order phase transition (i.e., proximity to ES).

Previously, we found evidence of ES in the brain networks of individuals experiencing abrupt wakefulness under light anesthesia^36^. Anesthesia reconfigures functional brain networks, decreasing global efficiency and enhancing modularity, which suppresses network synchronization and prompts ES conditions in some subjects^36,37^. Our EEG analysis showed that ES proximity in a resting state is significantly correlated with both induction and recovery times associated with propofol anesthesia, especially in the alpha band (8–13 Hz). Alpha waves coordinate hierarchical neural activities, facilitating cognitive processing and top-down control^38,39^, filter out irrelevant sensory inputs, enhance attentional focus, and transfer information globally via traveling waves^40,41^.

Our EEG analysis showed that only the alpha network’s ES proximity in the resting state reliably predicts both induction and recovery times. This finding suggests that the network mechanism driving diverse brain state transitions at the edge of consciousness prominently operates within globally networked alpha waves. Given that ES proximity significantly influences the brain’s response to perturbations, we propose that individual variations in a brain network’s proximity to ES should be considered an important factor for effectively predicting and modulating brain states through various brain stimulations (pharmacological, electrical, magnetic, etc.).

We applied the same method to financial networks during an economic crisis to test its generality. Significant correlations were found between the kurtosis of ACF values in the pre-crisis period, the market response time during the crisis, and the recovery time afterward. These results aligned with both our computational model predictions and observations related to anesthetic state transitions, suggesting that ES proximity, as measured by daily stock prices, could serve as a market index for characterizing financial markets and potentially forecasting market collapse and recovery. Furthermore, we discovered that ES proximity in the pre-crisis period is negatively correlated with a country’s GDP per capita, indicating that markets with higher ES proximity tend to have lower GDP. This implies that emerging markets with lower GDP per capita are closer to a first-order phase transition and are more vulnerable to economic crises than mature markets. Our findings underscore the importance of further research into the relationship between stock market network dynamics, GDP, and market collapse and recovery during crises. These results provide new insights into understanding economic crises through the lens of financial network dynamics and structure. Additionally, it remains unknown how these findings may apply to other types of markets—such as bonds, foreign exchange, derivatives, and cryptocurrency—and to different sources of economic crisis, such as the COVID-19 pandemic.

### Novel insights into rapid and prolonged system collapse and recovery around the time of crisis

First, research on catastrophic phase transitions, commonly known as “fold bifurcations” or “subcritical bifurcations” in bifurcation theory, has made significant theoretical progress^18-20,42^. Empirical indicators like critical slowing down, increasing variance and correlation in space and time, and spectral reddening (shifting the highest frequency to a lower one) have demonstrated the potential to anticipate impending phase transitions in diverse systems, including socioecological, neurological, financial, and climate systems. Recent theoretical studies have highlighted the need not only to predict upcoming critical points but also to identify the type of bifurcation that governs the transition pattern near these critical points. For instance, a smooth transition (transcritical bifurcation), the emergence of oscillations (Hopf bifurcation), or an abrupt change to another attractor (fold bifurcation) could characterize the transition pattern. However, given the generality of the critical slowing-down phenomenon, these indicators are unable to distinguish the bifurcation types, limiting the capacity to estimate the transition patterns near critical points^43^. In complex dynamical networks, the kurtosis of ACF offers a promising approach to address this limitation by estimating the proximity of a network’s phase transition type to a first-order transition, i.e., ES. We propose that combining the kurtosis of ACF with critical slowing-down indicators may enable the prediction of both impending critical points (abrupt transitions, often associated with system crises) and their associated phase transition types.

Second, previous theoretical models have identified network configurations that can induce ES^13^. However, practical application is limited by the difficulty of obtaining precise network structures in real-world scenarios. The kurtosis of ACF, a signal-based ES proximity indicator, can estimate ES proximity using time series signals, making it valuable for monitoring changes in a network’s phase transition type over time. For instance, we applied this approach to sickle cell disease and found that ES proximity in brain networks progressively increases until a pain crisis occurs and then diminishes afterward, repeating unpredictably over weekly or monthly intervals^44^. The kurtosis of ACF could be used to monitor changes in ES proximity and assess network vulnerability over time.

Third, our findings suggest the possibility of controlling abnormal network recoveries by modulating ES proximity. In previous modeling, we demonstrated that network modulation, specifically enhancing hub connections, converts the type of phase transition in the brain network from ES to non-ES, reducing sensitivity to external stimuli^16^. Another study showed that adding or removing a few key links can induce abrupt transition, called “ES bombs,” supporting the potential of modulating ES proximity through local structural changes^15^. This line of research could advance novel network modulation methods that could, for example, reduce the hypersensitivity in the brain in chronic pain, facilitate the recovery of normal brain functions in pathologic states, and accelerate market recovery after an economic crisis.

This study has several limitations. First, while the observed relationship between signal characteristics (kurtosis of ACF) at critical points, ES proximity, and critical state transitions is intriguing, it lacks analytical justification. Future research should aim to develop a mathematical framework to better understand this relationship. Second, the computational modeling used the maximal PCF of the instantaneous order parameter as a proxy for identifying the critical point. However, while a high PCF is necessary for criticality, it is not sufficient to confirm it. Due to this limitation, we prioritized evaluating the state transition rate after perturbation rather than performing advanced statistical tests to definitively verify criticality. Third, given the complexity of the study, we employed a simplified perturbation model to explore the influence of ES proximity on criticality loss and recovery rates. For feasibility, we assumed a uniform perturbation across all networks, because accounting for diverse perturbation forms was beyond the scope of this work.

Our study demonstrates that a network’s proximity to ES at a critical point plays a pivotal role in its resilience to external disruption. Specifically, the network’s susceptibility dictates the collapsing process, while the recovery is influenced by the internal resistance determined by ES proximity. This finding paves the way for developing network-specific methods to predict and modulate network collapse and recovery rates. This approach could be applied to various complex networks, including brain and financial networks, to enhance resilience and prevent or mitigate abrupt crises.

## Acknowledgment

This work was supported by the National Institute of General Medical Sciences (NIGMS) grant R21GM143521 to U.L. and D.P., the National Research Foundation of Korea (NRF) grant NRF-2022R1F1A1068796, the Institute of Information & Communications Technology Planning & Evaluation (IITP) grant No. 2022-0-00857, and the Ministry of Science and ICT (MSIT) grant No. 2022-0-00857, awarded to G.O.

## Author contribution

U.C., H.K., M.K., G.O., and G.M. conceived and designed the research project. H.K. and P.J. performed the computational modeling study. H.K. and M.K. analyzed the EEG data, which was curated by C.W. A.P. analyzed the stock data. U.C., D.P., and G.O. secured the funding. I.T., J.S., and G.M. provided critical revisions and feedback. U.C., H.K., G.O., A.P., D.P., and G.M. wrote and edited the manuscript. All authors discussed the results and approved the final version of the manuscript.

