## Supplementary material for "Proximity to Explosive Synchronization Determines Network Collapse and Recovery Trajectories in Neural and Economic Crises": Supplmentary table, figures, method

### Supplementary Information

#### Contents


- **S1. Computational Model**
  - Modified Stuart-Landau model with two new control parameters: one for modulating ES proximity and the other for stimulation strength
  - Changes in Kurtosis of ACF (KACF) and Kurtosis of PCF (KPCF) with varying adaptive feedback process strength (Z) (**Figure S1**) - (Statistical analyses: **Table S1** and **S2**)
  - Robustness tests with different stimulation strengths and durations (**Figure S2**)
    - Stimulation strengths: u(t)= 20, 40, 60, 80,100
    - Stimulation duration: 0.1, 0.5, 1, 2, 5,10, 20 seconds
  - Robustness tests with different network topologies (**Figure S3**)
    - Random, scale-free, and small-world networks
- **S2. Application to Human Brain Networks during Anesthesia**
  - Acquisition of human EEG data during anesthesia
  - EEG data preprocessing, KACF calculation, and statistical analysis
- **S3. Application to Stock Market Networks during Economic Crisis**
  - Acquisition of stock market data during the 2008 economic crisis (**Table S3**)
  - Stock data preprocessing, KACF and KPCF calculations, and statistical analysis
  - Robustness tests with varying baseline periods (**Table S4**)
  - Robustness tests with varying time periods (α) (**Table S5**)
  - Association of GDP per capita with ES proximity of stock market network (**Table S6**)


#### S1. Computational Model

##### Modified Stuart-Landau Model with Adaptive Feedback and Stimulation Terms:

A large-scale brain network model using coupled Stuart-Landau oscillators at criticality has successfully replicated the brain dynamics in a conscious resting state observed empirically with fMRI, MEG, and EEG[1-3]. Building on the approach, we developed a modified Stuart-Landau model to simulate large variances in state transitions, including the hysteresis phenomenon observed during general anesthesia[4]. These transitions are modulated at critical points, with varying degrees of Explosive Synchronization (ES) proximity. With the addition of a new stimulation term, we examined the relationship between the ES proximity, the loss and recovery of critical state under external perturbation, and the influence of network topology.

The adaptive feedback term $R_j^Z$ represents a recursive interaction process between a node and the other nodes in (S1):

$$\dot{z}_j(t) = \left\{\lambda_j + i\omega_j - \left|z_j(t)\right|^2\right\} z_j(t) + R_j^Z S \sum_{k=1}^{N} A_{jk} K_{jk} z_j\left(t - \tau_{jk}\right) + \beta\xi_j(t)), \quad j = 1,2,\dots,N \quad \text{(S1)}$$

Here, the state of the $j_{th}$ oscillator is determined by a complex variable $z_j = r_j\, e^{i\theta_j}$ at time *t*, where $|z_j|^2 = r_j^2$ and $r_j$ and $\theta_j$ are the amplitude and phase variables of the oscillator *j* at time *t*, respectively. $\omega_j$ is the natural frequency; $\lambda_j$ controls the decay rate of the amplitude; $S$ is a coupling strength among the oscillators; and $A_{jk}$ is the anatomical connection between node $j$ and $k$. $\xi_j(t)$ is a Gaussian white noise for each node *j* and added to the dynamics with the standard deviation β=0.05. The connection matrix $A$ varies with the topologies of the network models used (random, scale-free, small-world, and brain networks). For the brain network, it consists of 78 cortical regions constructed from average diffusion tensor imaging (DTI) of 80 young adults[5]. We set $A_{jk} = 1$, if a connection exists between node $j$ and $k$, and $A_{jk} = 0$ otherwise. $\tau_{jk}$ is a time delay between node $j$ and $k$, which is crucial in the human brain network model.

$R_j$ is defined as:

$$R_j = \frac{1}{2}\left(e^{i\theta_j} + \frac{1}{N}\sum_{K=1}^{N} e^{i\theta_k}\right) \quad \text{(S2)}$$

which represents the extent of the phase synchrony of node $j$ with the other nodes.

**The Control Parameter of ES Proximity Z:**

$Z$ is a scale term for the feedback process between one node and its connected nodes. Since $R_j$ is less than 1, a larger Z suppresses the interaction term with a smaller $R_j^Z$ in (S1) and pushes the network's phase transition type closer to ES. $R_j^Z$ with a larger *Z* inhibits the merging of small synchronization clusters, delaying the formation of a giant synchronization cluster in a network. This delayed transition triggers an abrupt synchronization transition at a critical point[6].

**Model Parameter Setting:**

- Natural Frequencies ($\omega_j$): Gaussian distribution around 10 Hz with a standard deviation of 0.4 Hz.
- Decay Rate of Amplitude ($\lambda_j$): Set to 1 for all oscillators.
- Time Delay ($\tau_{jk}$): Proportional to the physical distances between nodes *j* and *k* with a propagation speed of 7 m/s[7].
- Coupling Strength (*S*): Adiabatically increased and decreased from 0 to 4 with a step size of 0.05.

- Adaptive Feedback Strength ($Z$): Set to {0,0.5,1,1.5,2,2.5,3} to simulate distant and close ES proximities in the network.

We numerically solved the differential equations using the Stratonovich-Heun method with 1,000 discretization steps. One hundred different initial frequency configurations were simulated for each parameter set, and the results were averaged over all configurations.

**Identification of Critical Points in Network Models:**

An increase in signal variance and autocorrelation function is characteristic of a system approaching a critical transition, a phenomenon known as *critical slowing down*, where the system takes longer to return to equilibrium after a disturbance[8]. The variance (PCF) and the autocorrelation function (ACF) of the order parameters also reach their maximum at a critical point[9,10,11]. We explored the parameter space to identify the coupling strength that maximizes the PCF of the order parameters, considering the peak as a critical point. We examined the network dynamics characteristics at the critical points for 'distant' and 'close' ES proximities (Z) and studied their distinct response behaviors to external perturbation.

Notably, in this study, we focused on criticality transitions in networks with either 'distant' or 'close' proximity to ES, rather than in networks directly exhibiting ES, as ES rarely occurs in real-world networks under normal conditions. This implies that most of the networks we examined undergo second-order transitions rather than first-order phase transitions. When simulating networks with 'close' ES proximity (e.g., Z=3), the discrete nature of first-order transitions made it challenging to pinpoint the exact coupling strength that produces ES. Therefore, for a first-order transition, we defined the critical point as the coupling strength just before the discrete transition and evaluated ES in the time domain.

Dynamic noise introduced in the model allows a network in an incoherent state just before ES to cross the critical point, enabling observation of a discrete transition from incoherence to a highly synchronized state in the time domain (Figure S1. D). This transition is marked by significant changes in network dynamics, reflected in extreme ACF and PCF values and increased the kurtoses in ACF and PCF values. For consistency, we defined the critical point in non-ES networks as the coupling strength just before the PCF peak. Given the fine binning of coupling strengths, the KACF and KPCF values in non-ES networks closely match those observed at the ACF or PCF peak.

**Varying ES Proximity and Time to Critical State Loss/Recovery Under Perturbation**

We directly measured the speed of state transitions against an external perturbation by inducing a global pulsatile stimulus to the network and quantifying the change in network dynamics at critical points. The global pulsatile stimulus *u(t)* was applied as follows:

$$\dot{z}_j(t) = \left\{\lambda_j + i\omega_j - \left|z_j(t)\right|^2\right\} z_j(t) + R_j^Z S \sum_{k=1}^{N} A_{jk} K_{jk} z_j\left(t - \tau_{jk}\right) + \beta\xi_j(t) + \boldsymbol{u}(\boldsymbol{t}),$$

$$j = 1,2,\dots,N. \quad \text{(S3)}$$

$$u(t) = \begin{cases} p, & t_1 < t < t_2 \\ 0, & otherwise \end{cases}$$

where $p$ is the intensity of the stimulus during the period $T = t_2 - t_1$. We tested various stimulus strengths p = 10, 20, 40, 60, 80, 100, and durations $T$ = 0.1, 0.2, 0.5,1, 2, 5, 10, and 20 seconds, which are strong and long enough to influence the transition patterns during stimulation[10]. The global stimulus was applied at random timings $t_1$ for each trial. We conducted 20 trials for each of the 100 initial frequency configurations, resulting in 2,000 stimulus applications across the network at critical points.

The time to lose the critical state was defined as the time from stimulus onset until the order parameter exceeded three times the pre-stimulus standard deviation (at critical points). The time to recovery was defined as the time after stimulus cessation until the order parameter returned below three times the pre-stimulus standard deviation.

**Order Parameter:** The complex order parameter at time $t$ is defined as:

$$z(t) = r(t)e^{i\psi(t)} = \frac{1}{N}\sum_{f=1}^{N} e^{i\theta(j,t)}, \qquad \text{(S4)}$$

Where $z(t)$ is a complex order parameter at time $t$. $\psi(t)$ is the average phase over multi-variate time series signals at a time $t$; and $\theta(j,t)$ is the phase of a signal $j$ at a time $t$. The absolute value $r(t) = |z(t)|$ is the instantaneous order parameter at a time $t$, which represents the average phase synchronization over all individual signals at time $t$. $r(t)$ equals 1 when all signals are fully synchronized with the same instantaneous phase and 0 when the signals are completely incoherent, with the instantaneous phases uniformly distributed in $[0,2\pi)$ at time $t$.

**Calculation of Kurtoses of ACF values (KACF) and PCF values (KPCF):** The KACF/KPCF was calculated using the multivariate signals generated at a critical point as follows:

1. Extract instantaneous phases by applying the Hilbert transform to multivariate signals.
2. Calculate the instantaneous order parameters, defined in the coupled Kuramoto oscillators model.
3. Apply the moving window method to the instantaneous order parameters.
4. Calculate the autocorrelation function (ACF) and the pair correlation function (PCF) for each window
5. Calculate the kurtoses of ACFs and PCFs over the windows.

**Autocorrelation Function (ACF) and Pair Correlation Function (PCF) of Order Parameters:**

For a given data, we applied the moving window method with a window size that satisfies the pseudo-stationarity of dynamics, moving with a half-size window overlap. For each data in this study, the simulated signals, EEG signals, and daily stock returns, set an appropriate window size. The $ACF_k(\tau)$ of the order parameter $r(t)$ for a window $k$ is calculated for lag $\tau$ as:

$$ACF_k(\tau) = \frac{\sum_{t=1}^{N-\tau} (r(t)-\mu_k)(r(t+\tau)-\mu_k)}{\sum_{t=1}^{N} (r(t)-\mu_k)^2} \qquad \text{(S5)}$$

Where $\mu_k$ is the mean of $r(t)$ within window *k*, and *N* is the total number of the order parameter samples in a window *k*. An appropriate time lag $\tau$ was selected for each type of signal (simulation, EEG, and stock price), and the robustness of its selection was tested with several time lags.

Finally, for a given sequence of $ACF_k(\tau)$, *k* =1, …, *M*, where *M* is the total number of windows. The sample kurtosis is calculated using:

$$Kurtosis\ of\ ACF = \frac{N\ \sum_{k=1}^{M} (ACF_k - \mu_{ACF})^4}{(\sum_{k=1}^{M} (ACF_k - \mu_{ACF})^2)^2} \qquad \text{(S6)}$$

where $\mu_{ACF}$ represents the mean of the sequence of ACFs over all windows. The kurtosis of the ACFs measures whether the ACFs of order parameters *r(t)* are heavy-tailed compared to the normal distribution. We expect that a network with close ES proximity likely exhibits typical signal properties of a first-order phase transition near a critical point, such as intermittent and bistable transition dynamics, which are observed as outliers, compared to a network with distant ES proximity.

In addition, the Pair Correlation Function (PCF)[11] was calculated using:

$$PCF = \langle Re^2[z(t)] \rangle_t - \langle Re[z(t)] \rangle_t^2 \qquad \text{(S7)}$$

where $Re\ (z\ (t))$ denotes the real part of the complex order parameter at time *t*. The kurtosis of PCF was calculated using the same process as for ACF, with PCF replacing ACF. We found that the kurtosis of PCF (KPCF) is significantly correlated with the times of critical state loss and recovery in daily stock prices during crises. However, we couldn't find significant correlations in human EEG during anesthesia. Therefore, we presented the KACF results in the main text.

**Characteristic Dynamics at Critical Points of Networks with Different ES Proximities (Z)**
For Z=0 (Figure S1.A), the network exhibits typical dynamics near and far from the critical point. As the network approaches the critical points, the variance of instantaneous order parameters gradually increases, reflected by increasing PDF values. Based on previous studies[2,3,4,10,11], we defined the maximum PCF as the critical point. Across networks with varying ES proximities (from Z=0 to 3), the increase in PCF up to the critical points remains consistent. As Z increases (e.g., Z=1 and 2, shown in Figure S1.B and C), the network began to display intermittent state transitions at critical points, alternating between extended periods of highly synchronized state and desynchronized state. At Z=3, the network dynamics become sharply bistable, shifting abruptly to a highly synchronized state at the critical point.

**Calculation of KACF and KPCF and Statistical Analysis:** The temporal dynamics of the order parameter vary with different Z values. Time series signals were generated at critical points, with 300 seconds of data collected after an initial transition period. Using a moving window method (10-second window, 5-second overlap), we calculated the ACF and PCF of instantaneous order parameters within each window. For each Z value, KACF and KPCF were calculated from 59 ACF and 59 PCF values, respectively. Each simulation was repeated 100 times with random initial conditions. The mean and standard error of KACF and KPCF at critical points are shown in Figure S1 (E) and (F), respectively. The results indicate that as network's phase transition type approaches ES, the KACF and KPCF of network dynamics at critical points significantly increase, supporting both indexes as indicators of ES proximity. Statistical analysis, conducted via one-way ANOVA with post-hoc LSC and Tukey-Kramer tests, further validates these findings.

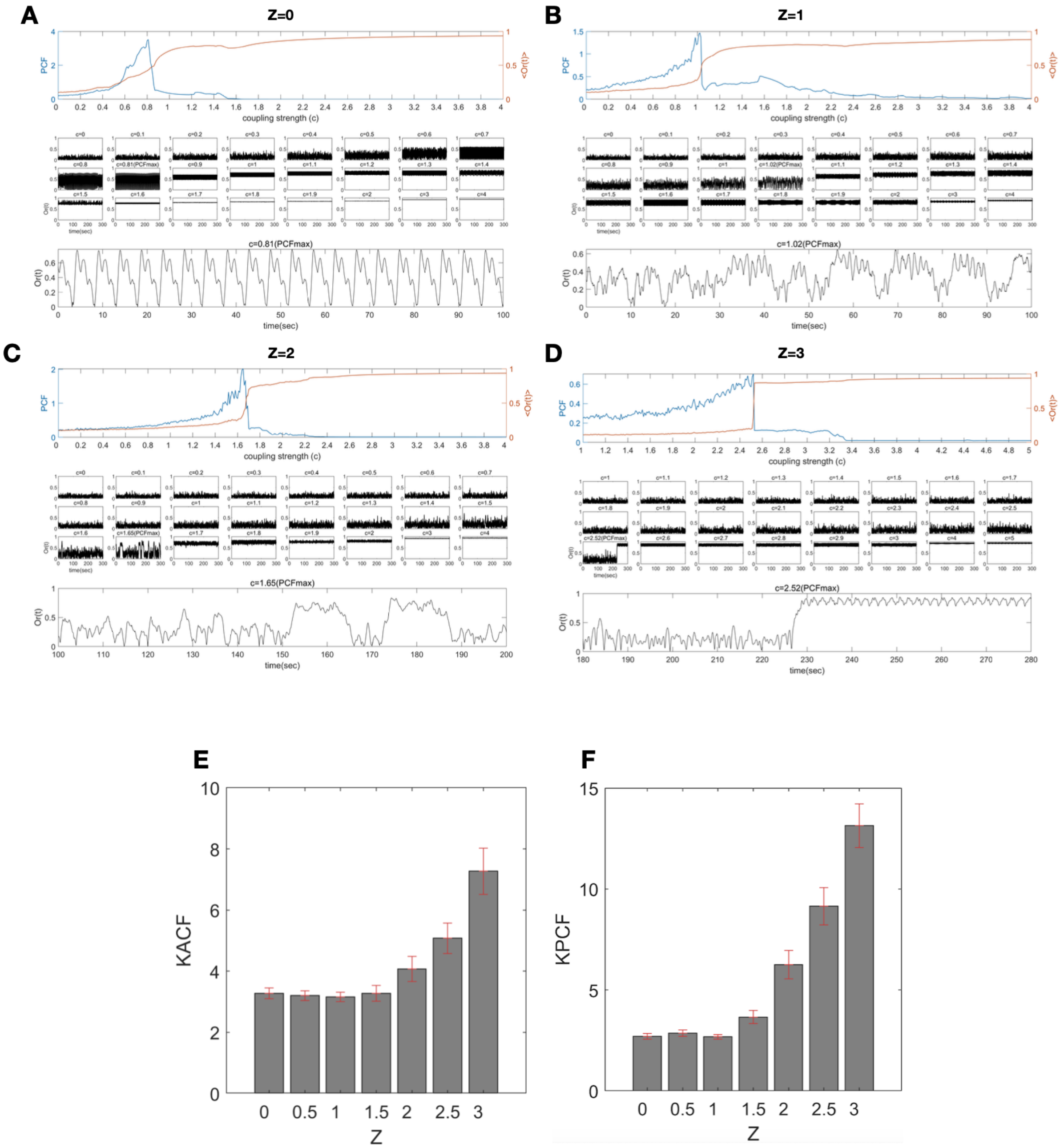


**Figure S1. KACF and KPCF of Instantaneous Order Parameters at Critical Points versus ES Proximity (Z) in Computational Models.**

Panels (A-D) provide examples illustrating changes in the PCF and the order parameters as coupling strength varies from 0 to 4, displaying characteristic network dynamics for ‘distant’ and ‘close’ ES proximities with adaptive feedback strengths (Z = 0, 1, 2, and 3). The first column shows how PCF changes with different coupling strengths and identifies the maximum PCF as the critical point for each network. The second column shows changes in the order parameter over time (300 seconds) for different coupling strengths. The third column presents fluctuations in the instantaneous order parameter near the critical point and quantifies its characteristic dynamics using KACF and KPCF. Panels (E) and (F) illustrate changes in KACF and KPCF as functions of ES proximity (Z) in the model, with error bars indicating standard errors.

**Statistical Analysis: Table S1**. Figure S1 (E), one-way ANOVA and post-hoc analysis, Multi-comparisons – 'LSD' and 'Tukey-Kramer' (Significant differences between groups are highlighted with gray): F(2,793) = 26.46.

**KACF**: Multi-comparison – 'LSD'

| Z | Z | difference | Lower 95% | Upper 95% | p-value |
|---|---|---|---|---|---|
| 0 | 0.5 | -0.644 | -0.143 | 0.358 | 0.577 |
| 0 | 1 | -0.780 | -0.280 | 0.221 | 0.275 |
| 0 | 1.5 | -0.963 | -0.463 | 0.0380 | 0.0705 |
| 0 | 2 | -1.107 | -0.606 | -0.105 | 0.0178 |
| 0 | 2.5 | -1.917 | -1.416 | -0.915 | 3.25e-08 |
| 0 | 3 | -3.119 | -2.619 | -2.118 | 3.14e-24 |
| 0.5 | 1 | -0.638 | -0.137 | 0.364 | 0.593 |
| 0.5 | 1.5 | -0.821 | -0.320 | 0.181 | 0.211 |
| 0.5 | 2 | -0.964 | -0.464 | 0.0370 | 0.069 |
| 0.5 | 2.5 | -1.774 | -1.274 | -0.773 | 6.58e-07 |
| 0.5 | 3 | -2.977 | -2.476 | -1.975 | 7.16e-22 |
| 1 | 1.5 | -0.684 | -0.184 | 0.317 | 0.473 |
| 1 | 2 | -0.828 | -0.327 | 0.174 | 0.201 |
| 1 | 2.5 | -1.638 | -1.137 | -0.636 | 8.91e-06 |
| 1 | 3 | -2.84 | -2.339 | -1.839 | 9.96e-20 |
| 1.5 | 2 | -0.645 | -0.144 | 0.357 | 0.574 |
| 1,5 | 2.5 | -1.455 | -0.954 | -0.453 | 0.000193 |
| 1.5 | 3 | -2.657 | -2.156 | -1.656 | 4.98e-17 |
| 2 | 2.5 | -1.311 | -0.810 | -0.310 | 0.00153 |
| 2 | 3 | -2.514 | -2.013 | -1.512 | 4.69e-15 |
| 2.5 | 3 | -1.704 | -1.203 | -0.702 | 2.63e-06 |

**KACF**: Multi-comparison - 'Tukey-Kramer'

| Z | Z | difference | Lower 95% | Upper 95% | p-value |
|---|---|---|---|---|---|
| 0 | 0.5 | -0.896 | -0.143 | 0.610 | 0.998 |
| 0 | 1 | -1.033 | -0.280 | 0.473 | 0.930 |
| 0 | 1.5 | -1.216 | -0.463 | 0.290 | 0.541 |
| 0 | 2 | -1.359 | -0.606 | 0.147 | 0.211 |
| 0 | 2.5 | -2.169 | -1.416 | -0.663 | 6.15e-07 |
| 0 | 3 | -3.372 | -2.619 | -1.865 | 0 |
| 0.5 | 1 | -0.890 | -0.137 | 0.616 | 0.998 |
| 0.5 | 1.5 | -1.073 | -0.320 | 0.433 | 0.874 |
| 0.5 | 2 | -1.217 | -0.464 | 0.289 | 0.539 |
| 0.5 | 2.5 | -2.027 | -1.274 | -0.521 | 1.28e-05 |
| 0.5 | 3 | -3.229 | -2.476 | -1.723 | 2.36e-22 |
| 1 | 1.5 | -0.937 | -0.184 | 0.569 | 0.992 |
| 1 | 2 | -1.08 | -0.327 | 0.426 | 0.862 |
| 1 | 2.5 | -1.89 | -1.137 | -0.384 | 0.000174 |
| 1 | 3 | -3.092 | -2.339 | -1.586 | 1.19e-19 |
| 1.5 | 2 | -0.897 | -0.144 | 0.609 | 0.998 |

| | | | | | |
|---|---|---|---|---|---|
| 1,5 | 2.5 | -1.707 | -0.954 | -0.201 | 0.00357 |
| 1.5 | 3 | -2.909 | -2.156 | -1.403 | 2.05e-16 |
| 2 | 2.5 | -1.563 | -0.810 | -0.0570 | 0.0255 |
| 2 | 3 | -2.766 | -2.013 | -1.26 | 3.63e-14 |
| 2.5 | 3 | -1.956 | -1.203 | -0.450 | 5.14e-05 |

**Statistical Analysis: Table S2.** Figure 1S (F), one-way ANOVA and post hoc analysis, $F(2,793) = 149.89$

##### **KPCF**: Multi-comparison – ‘LSD'

| Z | Z | difference | Lower 95% | Upper 95% | p-value |
|---|---|---|---|---|---|
| 0 | 0.5 | -1.280 | 0.07700 | 1.433 | 0.9110 |
| 0 | 1 | -1.329 | 0.02800 | 1.385 | 0.9670 |
| 0 | 1.5 | -2.101 | -0.7440 | 0.6120 | 0.2820 |
| *0* | *2* | -5.109 | -3.752 | -2.395 | 6.420e-08 |
| *0* | *2.5* | -9.930 | -8.573 | -7.216 | 2.380e-34 |
| *0* | *3* | -16.975 | -15.618 | -14.261 | 9.830e-104 |
| 0.5 | 1 | -1.406 | -0.04900 | 1.308 | 0.9440 |
| 0.5 | 1.5 | -2.178 | -0.8220 | 0.5350 | 0.2360 |
| *0.5* | *2* | *-5.186* | *-3.829* | *-2.472* | *3.450e-08* |
| *0.5* | *2.5* | -10.007 | -8.650 | -7.293 | 6.350e-35 |
| *0.5* | *3* | -17.052 | -15.695 | -14.338 | 1.160e-104 |
| 1 | 1.5 | -2.130 | -0.7730 | 0.5840 | 0.2640 |
| *1* | *2* | *-5.138* | *-3.781* | *-2.424* | *5.100e-08* |
| *1* | *2.5* | -9.959 | -8.602 | -7.245 | 1.450e-34 |
| *1* | *3* | -17.003 | -15.646 | -14.29 | 4.440e-104 |
| *1.5* | *2* | *-4.365* | *-3.008* | *-1.651* | *1.430e-05* |
| *1,5* | *2.5* | -9.186 | -7.829 | -6.472 | 4.800e-29 |
| *1.5* | *3* | -16.231 | -14.874 | -13.517 | 5.920e-95 |
| *2* | *2.5* | -6.179 | -4.822 | -3.465 | 4.030e-12 |
| *2* | *3* | -13.223 | -11.866 | -10.509 | 9.690e-63 |
| *2.5* | *3* | -8.402 | -7.045 | -5.688 | 6.350e-24 |

##### **KPCF**: Multi-comparison - 'Tukey-Kramer'

| Z | Z | difference | Lower 95% | Upper 95% | p-value |
|---|---|---|---|---|---|
| 0 | 0.5 | -1.964 | 0.07700 | 2.117 | 1 |
| 0 | 1 | -2.012 | 0.02800 | 2.069 | 1 |
| 0 | 1.5 | -2.785 | -0.7440 | 1.296 | 0.9360 |
| 0 | 2 | -5.792 | -3.752 | -1.712 | 1.230e-06 |
| 0 | 2.5 | -10.613 | -8.573 | -6.533 | 0 |
| 0 | 3 | -17.658 | -15.618 | -13.577 | 0 |
| 0.5 | 1 | -2.089 | -0.04900 | 1.991 | 1 |
| 0.5 | 1.5 | -2.862 | -0.8220 | 1.219 | 0.9000 |
| 0.5 | 2 | -5.869 | -3.829 | -1.789 | 6.530e-07 |
| 0.5 | 2.5 | -10.691 | -8.650 | -6.610 | 0 |
| 0.5 | 3 | -17.735 | -15.695 | -13.655 | 0 |
| 1 | 1.5 | -2.813 | -0.7730 | 1.267 | 0.9230 |
| 1 | 2 | -5.821 | -3.781 | -1.741 | 9.710e-07 |
| 1 | 2.5 | -10.642 | -8.602 | -6.562 | 0 |
| 1 | 3 | -17.687 | -15.646 | -13.606 | 0 |
| 1.5 | 2 | -5.048 | -3.008 | -0.9680 | 0.0002790 |
| 1,5 | 2.5 | -9.870 | -7.829 | -5.789 | 0 |
| 1.5 | 3 | -16.914 | -14.874 | -12.834 | 0 |
| 2 | 2.5 | -6.862 | -4.822 | -2.781 | 5.650e-11 |

| | | | | | |
|---|---|---|---|---|---|
| 2 | 3 | -13.906 | -11.866 | -9.826 | 0 |
| 2.5 | 3 | -9.085 | -7.045 | -5.005 | 0 |

#### Robustness of Simulation Results Across Different Stimulation Strengths and Durations

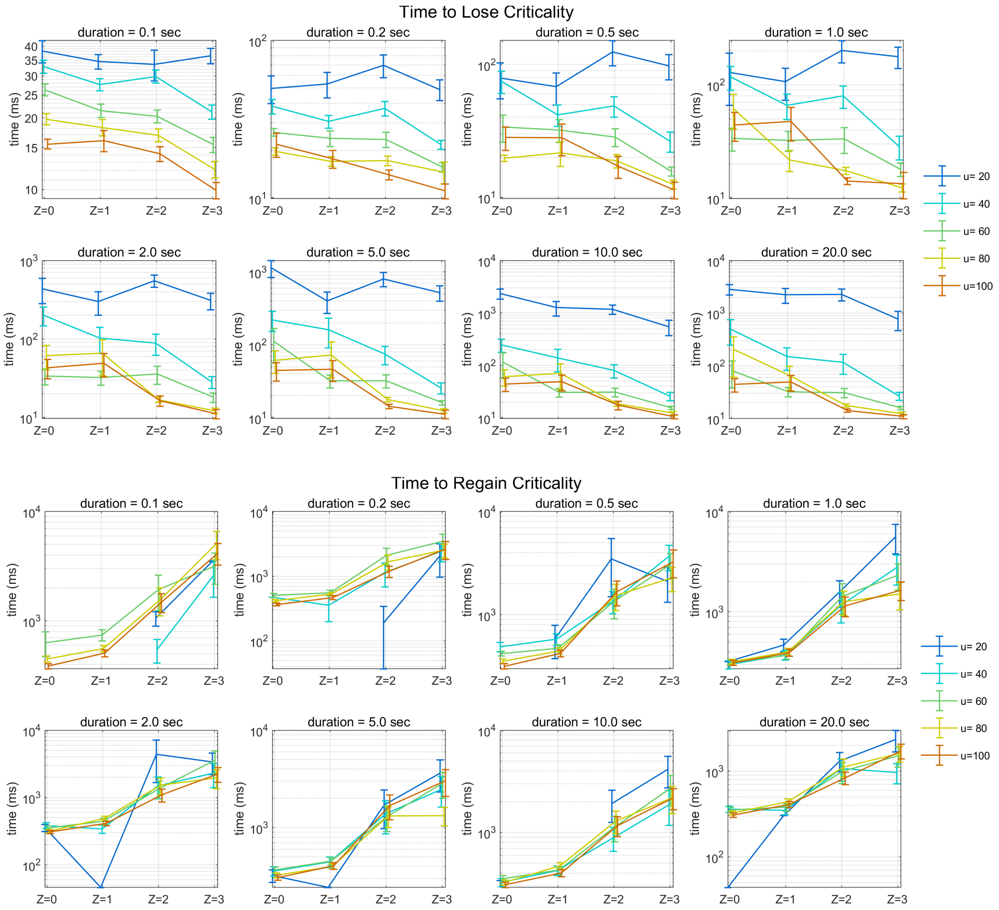


**Figure S2**. Relationships between ES proximity (adaptive feedback strength, Z), stimulation strength, stimulation duration, and the times to critical state loss and recovery. Networks with closer ES proximity (larger Z) exhibit shorter times to critical state loss and longer times to critical state recovery. This pattern is consistent across stimulation strengths ($u$=20, 40,60,80,100) and stimulation durations (0.1, 0.2, 0.5,1, 2, 5, 10, 20 seconds), except in cases where the stimulation is too weak ($u$=20) to deviate from the baseline dynamics. The relationship between ES proximity ($Z$) and the times to critical state loss and recovery becomes salient under external stimulations strong enough to push the network dynamics away from the critical state (u>20). Notably, the time scales of critical state loss and recovery depend on both the stimulation strength and duration.

**Robustness Tests with Different Network Topologies:** We tested the robustness of the simulation results using different network topologies including random, scale-free, small-world networks.

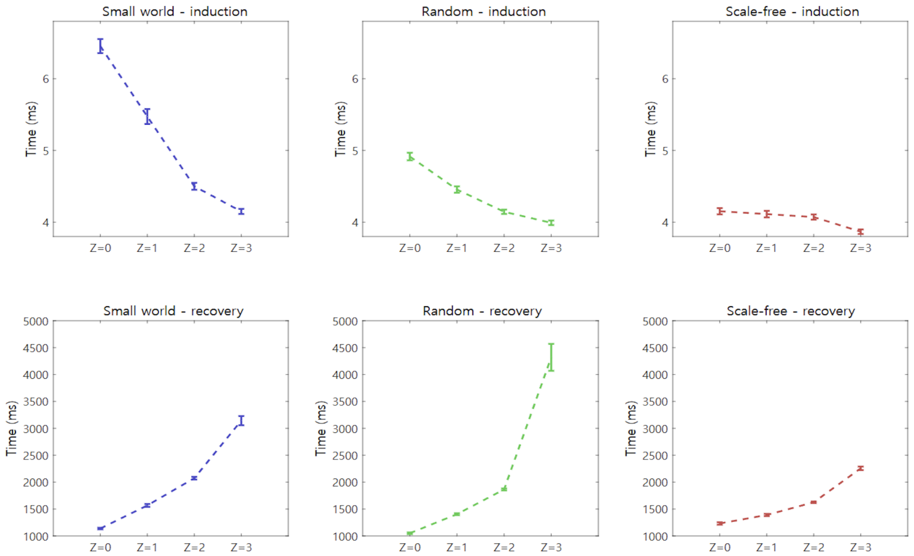


**Figure S3**. Relationships between ES Proximity (Adaptive Feedback Strength, Z), Network Topology, and Times to Critical State Loss and Recovery.

This figure shows how ES proximity (Z) relates to network topology and the time to critical state loss and recovery. Networks with closer ES proximity (larger Z) exhibit faster critical state loss and slower recovery, compared to networks with distant ES proximity (smaller Z). This trend is observed across all network topologies: (A) Small-world, (B) Random, and (C) Scale-free networks. Each network comprises 1,000 coupled Stuart-Landau oscillators (nodes) with an average node degree of 2. Error bars represent the standard errors from 100 simulations, each with different initial conditions. Both the small-world (Watts-Strogatz) network[12] and random (Erdős-Rényi) network[13] show higher sensitivity to changes in Z, with notable differences in the time to critical state loss and recovery. In contrast, the scale-free (Barabási-Albert) network[14] displays less sensitivity to variations in Z. This differential sensitivity is due to the characteristic topologies of these networks. The high clustering in small-world networks and the heterogeneity in random networks may enhance conditions for ES, inhibiting the formation of a giant synchronized cluster even at the same feedback strength (Z). On the other hand, the presence of highly connected hubs in scale-free networks may facilitate smoother phase transitions, reducing the likelihood of delayed transitions. These results suggest that while the relationship between ES proximity and critical state loss and recovery is maintained across different topologies, the nature of the phase transition at the critical state is influenced by network topology.

#### S2. Application to Human Brain Networks during Anesthesia

**EEG Experimental Procedures:** We utilized 32-channel EEG data from 16 healthy subjects (8 male, 8 female; age 28.6±7.028.6±7.0 years; range 19–43 years) participating in a study involving an ultraslow target-controlled intravenous infusion of propofol, a γ-aminobutyric acid type A ($GABA_A$) agonist. The experiment was designed to observe brain dynamics during the induction of and emergence from deep sedation induced by propofol[15]. The central nervous system propofol effect-site concentration (ESC) was estimated throughout the experiment.

The EEG data comprised four distinct phases:

1. **Baseline Resting State**: 10 minutes of eyes-closed resting state to establish baseline measurements.
2. **Induction Phase**: Ultraslow propofol-induced loss of consciousness over 48 minutes, during which the estimated ESC gradually increased to a maximum of 4 μg/ml.
3. **Maintenance Phase**: A 10-minute period maintaining the peak propofol dose to observe sustained effects on brain activity.
4. **Recovery Phase**: Observation of brain activity during the recovery of consciousness after the propofol infusion was discontinued.

During phases 2 and 4, laser stimuli, auditory stimuli, and cognitive word tasks were randomly delivered at 1-minute intervals to assess sensory processing and cognitive function. The Loss of Behavioral Response (LOBR) was defined as the time point when a subject ceased responding to the word task, indicating a transition into unconsciousness. Conversely, the Recovery of Behavioral Response (ROBR) was marked by the time point when the subject resumed responding, indicating a return to consciousness. These data have been previously analyzed and published in separate studies[15,16]. Detailed experimental designs and protocols are thoroughly described in these prior publications.

**EEG Preprocessing**: EEG data were re-referenced using the average reference method and downsampled to 125Hz from 500Hz. A windowed sinc-FIR filter (EEGLAB MATLAB toolbox) was applied to prevent phase shifting in the signals. A bandpass filter was applied to isolate the alpha-frequency band (8–13 Hz). The Hilbert transform was then used to extract instantaneous phases from 31-channel EEG signals (excluding one reference channel) and calculated the instantaneous order parameters using (S4).

**KACF Calculation**: The moving window method was employed with a window size of 10 seconds and a 5-second overlap between windows. ACF values were calculated for each window using a time lag of 11, reflecting the dynamics of the alpha band, ~10Hz (as in equation S4). The KACF was computed from the ACF values across all windows.

**Statistical Analysis**: The Spearman correlation coefficient was calculated between the KACF values and the induction and recovery times of 16 subjects.

##### S3. Application to stock market networks during economic crisis:

This supplementary information on economic data analysis provides further details on the methods used to study financial market crises in 39 countries. It outlines the construction of the financial data sample, including the selection criteria for firms in each country, and provides a detailed explanation of the data analysis procedures. The section also addresses the robustness of the results, focusing on the choice of the baseline period and the time window used to identify market collapse and recovery during the economic crisis. Lastly, it explores the relationship between GDP per capita and the ES proximity of financial networks.

| ISO | COUNTRY | Market Index | #Stocks | Start(S) | End(E) | Max | Min | Emerging market | GDP |
|---|---|---|---|---|---|---|---|---|---|
| GBR | UK | FTSE 100 | 886 | 20080101 | 20090631 | 20070615 | 20090303 | | 44590 |
| THA | THAILAND | SET Index | 231 | 20071101 | 20090531 | 20071029 | 20081029 | O | 3370 |
| MEX | MEXICO | S&P_BMV IPC | 46 | 20080601 | 20090531 | 20071018 | 20081027 | O | 9457 |
| TUR | TURKEY | BIST 100 | 276 | 20080101 | 20090331 | 20071015 | 20081120 | O | 7961 |
| FIN | FINLAND | OMX Helsinki 25 | 93 | 20080201 | 20090631 | 20070713 | 20090309 | | 41309 |
| NLD | NETHERLANDS | AEX | 129 | 20080501 | 20090601 | 20070716 | 20090309 | | 44936 |
| CHE | SWITZERLAND | SMI | 165 | 20080501 | 20090531 | 20070601 | 20090309 | | 59221 |
| DEU | GERMANY | DAX | 347 | 20080301 | 20090631 | 20070716 | 20090306 | | 36894 |
| IDN | INDONESIA | IDX Composite | 70 | 20080501 | 20090531 | 20080109 | 20081028 | O | 1766 |
| HUN | HUNGARY | Budapest SE | 16 | 20080601 | 20090631 | 20070723 | 20090312 | O | 11483 |
| POL | POLAND | WIG20 | 147 | 20080301 | 20090731 | 20071029 | 20090217 | O | 9032 |
| PAK | PAKISTAN | Karachi 100 | 42 | 20071101 | 20090531 | 20080418 | 20090126 | O | 1032 |
| MYS | MALAYSIA | KLCI | 493 | 20071101 | 20090531 | 20080111 | 20081028 | O | 6351 |
| EGY | EGYPT | EGX 30 | 53 | 20071101 | 20090531 | 20080505 | 20090205 | O | 1564 |
| BEL | BELGIUM | BEL 20 | 95 | 20080301 | 20090431 | 20070523 | 20090306 | | 38842 |
| FRA | FRANCE | CAC 40 | 444 | 20080201 | 20090631 | 20070601 | 20090309 | | 37796 |
| IRL | IRELAND | ISEQ Overall | 23 | 20071201 | 20090831 | 20070601 | 20090309 | | 53739 |
| SAU | SAUDI ARABIA | Tadawul All Share | 51 | 20071101 | 20090531 | 20080112 | 20090309 | O | 17827 |
| AUS | AUSTRALIA | S&P_ASX 200 | 573 | 20080201 | 20110231 | 20071101 | 20090306 | | 40675 |
| ARG | ARGENTINA | S&P Merval | 24 | 20071101 | 20090531 | 20071031 | 20081121 | O | 5976 |
| CHL | CHILE | S&P CLX IPSA | 36 | 20080301 | 20090701 | 20070703 | 20081010 | O | 9410 |
| AUT | AUSTRIA | ATX | 45 | 20080301 | 20090731 | 20070709 | 20090309 | | 40675 |
| HKG | HONG KONG | Hang Seng | 129 | 20071101 | 20090531 | 20071030 | 20081027 | | 28028 |
| NZL | NEWZEALAND | NZX 50 | 47 | 20071101 | 20090531 | 20071002 | 20090303 | | 26220 |
| NOR | NORWAY | Oslo OBX | 90 | 20071101 | 20100831 | 20070719 | 20081121 | | 74256 |
| PRT | PORTUGAL | PSI 20 | 30 | 20080301 | 20090431 | 20070723 | 20090303 | | 19840 |
| ESP | SPAIN | IBEX 35 | 114 | 20080301 | 20090831 | 20071108 | 20090309 | | 28414 |
| KOR | KOREA | KOSPI | 936 | 20080201 | 20090531 | 20071031 | 20081024 | O | 21731 |
| IND | INDIA | Nifty 50 | 729 | 20081001 | 20090331 | 20080108 | 20081027 | O | 802 |
| JPN | JAPAN | Nikkei 225 | 2664 | 20080301 | 20090431 | 20070709 | 20090310 | | 36022 |
| RUS | RUSSIA | MOEX Russia | 12 | 20071101 | 20090531 | 20071212 | 20081024 | O | 7416 |
| CHN | CHINA | Shanghai Composite | 162 | 20080101 | 20090231 | 20071016 | 20081104 | O | 2095 |
| USA | USA | S&P 500 | 1723 | 20071101 | 20090531 | 20071009 | 20090309 | | 46217 |
| ITA | ITALY | FTSE MIB | 229 | 20080301 | 20090531 | 20070518 | 20090309 | | 33448 |
| PHL | PHILIPPINES | PSEI Composite | 47 | 20071101 | 20090531 | 20071008 | 20081028 | O | 1471 |
| SGP | SINGAPORE | STI | 217 | 20071101 | 20090531 | 20071011 | 20090310 | | 33768 |
| TWN | TAIWAN | TWII | 820 | 20071101 | 20090531 | 20071030 | 20081120 | O | 16893 |
| CAN | CANADA | S&P_TSX | 1070 | 20070901 | 20090731 | 20070719 | 20090309 | | 40559 |
| BRA | BRAZIL | Bovespa | 75 | 20080601 | 20090331 | 20080520 | 20081027 | O | 6067 |

**Table S3**. Details of economic crises in 39 countries. This table provides detailed information regarding the stock market and recession periods for 39 countries analyzed in this study. The first and second columns contain the ISO country codes and full names of the countries, respectively. The third and fourth columns specify the benchmark stock index used for analysis in each country and the number of companies analyzed from each country's stock market. The fifth and sixth columns indicate the start and end dates of the recession periods for each country, as per the OECD recession indicator. For the countries without the recession indicator, the U.S. recession period is used as a reference. The seventh and eighth columns provide the dates when the stock markets of each country reached their maximum and minimum values during the analysis period, capturing the peaks and troughs in market performance. The last two columns represent the emerging market indicator and the GDP per capita (US dollar) for each country from the International Monetary Fund in 2006.

**Construction of Stock Market Data:** Financial market data for this study were sourced from S&P Compustat Global and the Center for Research in Security Prices (CRSP) databases. These comprehensive databases provide detailed information on a wide range of global firms, including daily stock prices, shares outstanding, total assets, trading volume, exchange codes, and share codes. The number of firms analyzed varied between 11 and 2,664, depending on the country. This dataset provided the foundation for examining the economic impact of the 2008 global financial crisis across different countries.

The countries included were selected based on the availability of high-quality stock market data, representing a mix of developed and emerging economies to capture a wide range of market dynamics. Table S3 provides key details such as the representative stock market index for each country, the number of companies included, recession periods, and GDP per capita. For countries without OECD recession indicators (e.g., Thailand, Pakistan, Malaysia), the U.S. recession period was used as a reference.

**Data Selection Criteria:**

- **Firm-Level Data**: We focused on common stocks (share codes 10 and 11), which typically represent ordinary shares listed on the primary exchange in each country. Companies that reported negative total assets at any time during the analysis period were excluded to ensure data integrity. In cases where multiple share classes existed, the common share class with the highest total stock assets was chosen for consistency.
- **Country-Level Criteria**: Countries with insufficient data or irregular recession periods were excluded. For instance, Denmark was excluded because its recession began too early (July 2006), while Greece was excluded because its economic recovery extended beyond the study's timeframe (until 2011). After these exclusions, 39 countries remained, categorized as either developed or emerging markets, allowing for comparative analysis across different economic contexts.

**Dataset Overview:** The dataset was carefully curated to include diverse economies and geographic regions, aiming to analyze the heterogeneous impact of the 2008 financial crisis. Countries were selected based on the availability of sufficient stock data, focusing on both developed and emerging economies to represent different stages of economic development.

- **Stock Data Sources**: We collected daily stock data from the main exchanges in each country. In cases where a country had multiple significant exchanges, data from all relevant exchanges were included to ensure a representative sample size.
- **U.S. Companies**: Classified using the Permno code from the Center for Research in Security Prices (CRSP) database. The focus was on companies listed on the New York Stock Exchange (NYSE) and the American Stock Exchange (AMEX).
- **Other Countries**:
- For South Korea, both KOSPI and KOSDAQ were included.
- For India, companies from both the National Stock Exchange (NSE) and the Bombay Stock Exchange (BSE) were considered.
- For Russia, we included companies from the Moscow Exchange (MOEX), the Russian Trading System (RTS), and the Saint Petersburg Exchange.
- For China, firms listed on the Shanghai Stock Exchange (SSE) and the Shenzhen Stock Exchange (SZSE) were analyzed.
- For Taiwan, companies listed on both the Taiwan Stock Exchange (TWSE) and the Taipei Exchange (TPEx) were included.

**Pre-Crisis Normal Period:** The **"normal period"** was defined as January 2006 to December 2006, chosen as a stable phase before the onset of the 2007-2009 global financial crisis. This period served as a baseline to evaluate the ES proximity of each country's stock market under normal, pre-crisis conditions.

- **Data Cleaning**: During the sample period from July 2005 to December 2006, we removed companies with more than 95% zero values for their stock prices. This ensured that only actively traded firms were included, enhancing the reliability of the analysis.
- **ES proximity Measure**: To estimate KACF, we used a window size of 120 days. Figures 5C and 5D present the Spearman correlations between the KACF and response and recovery times, demonstrating the linkage between market conditions during the pre-crisis period and subsequent market performance during the crisis.
- **Sensitivity Analysis**: To ensure the robustness of our findings, we conducted a sensitivity analysis by systematically varying the start point of the pre-crisis period. We adjusted the starting point within a six-month window (from July 2005 to January 2006) and recalculated the ES proximity (KACF) for each country. The results were consistent, confirming the significant relationship between ES proximity across different pre-crisis periods and market response/recovery times during the crisis (as shown in Table S4).

**Economic Data Analysis Procedure:** In this study, we define a country's market collapse and recovery time during an economic crisis using changes in the stock market index (e.g., S&P500

index), while individual firm stock prices (e.g., Apple Inc.) are utilized to calculate market dynamics such as ES proximity or market criticality. A stock market index is a statistical measure reflecting the overall performance of a specific group of stocks (e.g., the S&P 500 index, which tracks 500 large US companies; the Nasdaq, focused on technology stocks). The stock market index $P_c^m$ of a country c and the stock price $P_c^f$ of a firm *f* are analogous to anesthetic concentration and a single-channel EEG signal in the brain, respectively.

**Definition of response time and recovery time:** The stock return $R_c^f(t)$ of a firm (*f*) in 39 countries (*c*) is computed as the difference of the natural logarithms of adjusted stock prices.

$$R_c^f(t) = log\ (aP_c^f(t)) - log\ (aP_c^f(t-1)\ ),\ \{f = 1,2,\dots,N_c\} \in SC_c$$

$$aP_c^f(t) = \left(\frac{P_c^f(t)}{ajexdi^f(t)}\right) \times Trfd^f(t) \qquad \text{(S8)}$$

Where $SC_c$ means the set of available firms *f* in a country $c$, and $t$ represents the daily time frequency from July 2005 to December 2006. To measure the stock returns $R_c^f(t)$, we first calculated the adjusted stock prices $aP_c^f(i,t)$ for all available firms of each country by using the information about $P_c^f(i,t)$, $ajexdi(i,t)$, and $Trfd(i,t)$ from S&P Compustat Global. $P_c^f(t)$ means the closing stock price of a firm $f$ listed in major exchange of country $c$. $ajexdi(i,t)$ is adjusted factor for all stock splits and dividends, providing a more accurate reflection of a company's equity value over time by removing the effects of these corporate actions of firm $i$ at time $t$ and Trfd is a daily factor that captures the total return of a stock, encompassing both price appreciation and dividend reinvestment, providing a comprehensive measure of investment performance. This price adjustment ensures that the stock returns reflect the effects of corporate actions, such as stock splits, mergers, acquisitions, and dividends.

We tested whether the KACF measured during the normal period correlates with each country's response and recovery time in the 2008 economic crisis. To account for differences in stock market scales across 39 countries, we normalized stock market index changes during the crisis and defined market collapse and recovery rates based on the extent of each country's price drop, allowing direct comparisons. Response rate (R1) and recovery rate (R2) were measured using the equation (S2).

$$R1(c) = \frac{P_c^m(t_1) - P_c^m(t_1+\alpha)}{P_c^m(t_1) - P_c^m(t_2)}\ ,\ R2(c) = \frac{P_c^m(t_2+\alpha) - P_c^m(t_2)}{P_c^m(t_1) - P_c^m(t_2)} \qquad \text{(S9)}$$

Where $P_c^m(t)$ is the stock market index of country $c$ and time $\mathrm{t} = 1, 2, 3, \dots \mathrm{T}$. $t_1 = \{t \in\ T : \max P_c^m(t)\}$ and $t_2 = \{t \in T : \min P_c^m(t)\}$, indicate the dates of the maximal and minimal stock market index in country $c$ ,respectively. These rates mean the price difference between the maximal (minimum) price and a reference point, which is how much the stock market index decreases(increases) from the maximum (minimum) prices for a given time $\alpha$ (illustrated in Figure 5B). We assumed that the maximum and minimum prices for each country during the 2008 crisis indicate, respectively, the turning points of a contrary market to enter and escape its crisis.

We set the reference point as the price after $\alpha$ days from the maximal (minimum) price. Because the 39 countries show diverse market dynamics, we tested several time periods, $\alpha = \{40, 60, 80, 100,\ and\ 120\ days\}$. 40 and 120 days approximately correspond to 2 and 6 months, considering market closing days. We chose $\alpha$ of 100 days, an appropriate period to reflect the temporal scale of the overall price changes during the worldwide economic crisis. The sensitivity of the $\alpha$ selection was tested in Table S4 and 5. Finally, we applied a logarithm to the inverse of the market response and recovery rates for direct comparison with the EEG study's induction time and recovery time.

$$Response\ time(c) = \ln\left(\frac{1}{R1(c)}\right),\ Recovery\ time(c) = \ln\left(\frac{1}{R2(c)}\right), \qquad \text{(S10)}$$

If the country has a higher response (recovery) rate, it represents a fast financial market collapse (recovery). We anticipate the country's stock market will be healthy if it has a longer response time and a shorter recovery time.

**Indicators of ES proximity of a stock market: KACF and KPCF:** In this section, we investigate the hypothesis that the ES proximity of a stock market network may influence its resilience and recovery capabilities during the 2008 economic crisis. Our analysis focuses on the relationship between the ES proximity and the times to respond to and recover from the crisis. Using the same procedure as in the EEG study, we computed the KACF as follows: (1) extract the phases of stock returns, (2) calculate the ACFs of the moving window, (3) calculate the KACF over all the windows.

1. Apply the Hilbert transform to the daily stock returns $R_c^f(t)$ of the normal period from July 1st, 2005 to December 31st, 2006, and get the instantaneous phases $Z_c^f(t)$ for each firm *f* in a country *c*.
2. Calculate the instantaneous order parameters with the multivariate phases $Z_c^f(t)$ of listed firms for each country *c* using S4.
3. Calculate the ACF of each window (window size of 120 days, 1 day overlapping) using (S5), which produces about 250 ACF values during the normal period for each country. We tested different time lags ($\tau$ =1,2, and 3) and presented the robustness of the $\tau$ selection in Table S5. In the main text, we present the result of $\tau$= 2.
4. Calculate the KACF over the normal period for a county using (S6). Repeated this procedure (1), (2), and (3) for 39 countries.

In addition, we tested ES proximity using an alternative criticality indicator, Pair Correlation Function (PCF). We repeated the procedure (1)-(3) after replacing ACF with PCF, defined in (S7). We demonstrated that the kurtosis of PCF (KPCF) during the normal period has significant correlations with both the response time and the recovery time, as shown in Table S6.

Notably, despite KACF showing correlations in all the data (simulated data, EEG, and stock price), we did not find a significant correlation with KPCF in the EEG study. This might be influenced by the relatively small number of channels (31 channels). Considering that the order parameter measures the global phase coherence in a network, a small number of samples may be limited to reflect the global network dynamics of the brains in anesthesia.

**Robustness to the varying normal periods, alpha $\alpha$, and time lag:** In Table S5, we evaluated how the selection of the "normal period" influences the Spearman correlations between response/recovery times and the KACF. The normal period was defined based on the principles of the Efficient Market Hypothesis (EMH) to ensure it represents a stable phase before the onset of the subprime financial crisis. We tested three different normal periods and nine values of alpha (α), which represent the duration after the market's minimum (or maximum) point, used to calculate both response and recovery times. Our results indicate that the choice of the normal period does not significantly affect the correlation between market response/recovery times and the KACF.

In Table S5, we tested the robustness of the KACF correlation by varying both the time lags (1, 2, and 3) and the alpha (α) values (ranging from 40 to 120 days). We found that the KACF with a time lag of 2 showed a significant correlation with both response and recovery times. Additionally, testing with KPCF also revealed significant correlations with response/recovery times, but only for alpha (α) of 40 days. This suggests that KPCF is more sensitive to the specific methods used for defining response and recovery times.

**Relationship between GDP per capita and market ES proximity:** To test the robustness of the potential relationship between countries' economic development and the stock market's ES proximity, we analyzed Gross Domestic Product (GDP) per capita for 39 countries in 2006, selected as the pre-crisis year (see Table S3). We evaluated two ES proximity indicators - KPCF and KACF at lags 2- using various normal periods. Table S6 shows significant negative Spearman correlations between both ES proximity indicators and GDP per capita. This correlation persisted even when adjusting the normal period. The observed correlations between ES proximities of countries' stock market networks and the levels of economic development may explain why 'developed' countries, with more distant ES proximity, tend to be more resilient during the crisis compared to ' 'developing' countries, which exhibit closer ES proximity and are more susceptible to phase transitions.

**Table S4**. The robustness of varying baseline periods and α

| | | alpha(α) | | | | | | | | |
|---|---|---|---|---|---|---|---|---|---|---|
| | | 40 | 50 | 60 | 70 | 80 | 90 | 100 | 110 | 120 |
| Response time | 200511 | -0.08 | -0.12 | -0.23 | -0.36** | -0.40** | -0.43*** | -0.38** | -0.34** | -0.34** |
| (p-value) | | (0.63) | (0.47) | (0.15) | (0.03) | (0.01) | (0.01) | (0.02) | (0.03) | (0.03) |
| | 200509 | -0.11 | -0.16 | -0.27 | -0.38** | -0.43*** | -0.45*** | -0.40** | -0.37** | -0.37** |
| | | (0.49) | (0.33) | (0.10) | (0.02) | (0.01) | (0.00) | (0.01) | (0.02) | (0.02) |
| | 200507 | -0.10 | -0.14 | -0.25 | -0.37** | -0.41*** | -0.44*** | -0.39** | -0.35** | -0.35** |
| | | (0.56) | (0.40) | (0.13) | (0.02) | (0.01) | (0.01) | (0.02) | (0.03) | (0.03) |
| Recovery time | 200511 | 0.42*** | 0.36** | 0.41** | 0.31* | 0.37** | 0.51*** | 0.47*** | 0.40** | 0.40** |
| (p-value) | | (0.01) | (0.02) | (0.01) | (0.06) | (0.02) | (0.00) | (0.00) | (0.01) | (0.01) |
| | 200509 | 0.43*** | 0.35** | 0.40** | 0.28* | 0.36** | 0.50*** | 0.45*** | 0.37** | 0.37** |
| | | (0.01) | (0.03) | (0.01) | (0.09) | (0.03) | (0.00) | (0.00) | (0.02) | (0.02) |
| | 200507 | 0.42*** | 0.36** | 0.41** | 0.30* | 0.37** | 0.51*** | 0.47*** | 0.39** | 0.39** |
| | | (0.01) | (0.03) | (0.01) | (0.06) | (0.02) | (0.00) | (0.00) | (0.01) | (0.01) |

This table shows the Spearman correlations between response(recovery) time using different alpha ($\alpha$) and KACF2 estimated from the different normal periods. KACF2 means kurtosis of autocorrelation function with time lag 2. ***, **, and * represent significant levels at 1%, 5%, and 10%, respectively. Values in parentheses are t-statistics. The parameter $\alpha$ indicates the duration after the minimum (maximum) point of the stock market index, which is used to define both the response time and recovery time in Equation (S10). Three different baseline periods were tested.

**Table S5**. The robustness to varying time periods ($\alpha$)

| Criticality Indicators | alpha(α) | | | | | | | | |
|---|---|---|---|---|---|---|---|---|---|
| | 40 | 50 | 60 | 70 | 80 | 90 | 100 | 110 | 120 |
| Panel A: Response time (p-value) | | | | | | | | | |
| KPCF | -0.49*** | -0.45*** | -0.26 | -0.30* | -0.30* | -0.39** | -0.37** | -0.41** | -0.32** |
| | (0.00) | (0.00) | (0.11) | (0.07) | (0.06) | (0.02) | (0.02) | (0.01) | (0.04) |
| KACF1 | 0.15 | 0.09 | -0.04 | -0.07 | -0.14 | -0.06 | -0.06 | 0.02 | -0.08 |
| | (0.37) | (0.60) | (0.81) | (0.68) | (0.41) | (0.70) | (0.73) | (0.91) | (0.62) |
| KACF2 | -0.13 | -0.15 | -0.27 | -0.38** | -0.43*** | -0.45*** | -0.40** | -0.37** | -0.38** |
| | (0.45) | (0.36) | (0.10) | (0.02) | (0.01) | (0.00) | (0.01) | (0.02) | (0.02) |
| KACF3 | -0.28* | -0.21 | -0.28* | -0.45*** | -0.53*** | -0.53*** | -0.59*** | -0.54*** | -0.50*** |
| | (0.09) | (0.19) | (0.08) | (0.00) | (0.00) | (0.00) | (0.00) | (0.00) | (0.00) |
| Panel B: Recovery time (p-value) | | | | | | | | | |
| KPCF | 0.28* | 0.19 | 0.20 | -0.03 | 0.15 | 0.22 | 0.26 | 0.18 | 0.15 |
| | (0.08) | (0.25) | (0.23) | (0.83) | (0.35) | (0.19) | (0.12) | (0.27) | (0.36) |
| KACF1 | 0.07 | 0.02 | 0.16 | 0.03 | 0.17 | 0.24 | 0.10 | 0.13 | 0.07 |
| | (0.69) | (0.92) | (0.33) | (0.84) | (0.29) | (0.15) | (0.53) | (0.42) | (0.68) |
| KACF2 | 0.44*** | 0.37** | 0.40** | 0.28* | 0.35** | 0.50*** | 0.49*** | 0.42*** | 0.43*** |
| | (0.01) | (0.02) | (0.01) | (0.08) | (0.03) | (0.00) | (0.00) | (0.01) | (0.01) |
| KACF3 | 0.26 | 0.17 | 0.22 | 0.01 | 0.06 | 0.21 | 0.23 | 0.21 | 0.12 |
| | (0.12) | (0.30) | (0.19) | (0.96) | (0.72) | (0.19) | (0.16) | (0.21) | (0.47) |

This table shows the Spearman correlations between response(recovery) time using different alpha ($\alpha$) and KACF(KPCF). KACF1 means kurtosis of autocorrelation function with time lag 1. KPCF means kurtosis of the pair correlation function. ***, **, and * represent significant levels at 1%, 5%, and 10%, respectively. Values in parentheses are t-statistics. The parameter $\alpha$ indicates the duration after the minimum (maximum) point of the stock market index, measured at daily frequency, which is used to define both the response time and recovery time in Equation (S1).

**Table S6**. Spearman correlations between GDP and KACF(KPCF) with adjusting normal period

| | Normal period | | | | | | |
|---|---|---|---|---|---|---|---|
| ES | 200507 | 200508 | 200509 | 200510 | 200511 | 200512 | 200601 |
| proximity | ~ 200612 | ~ 200612 | ~ 200612 | ~ 200612 | ~ 200612 | ~ 200612 | ~ 200612 |
| KACF2 | -0.19 | -0.32** | -0.33** | -0.30* | -0.35** | -0.30* | -0.36** |
| (p-value) | (0.26) | (0.05) | (0.04) | (0.06) | (0.03) | (0.07) | (0.02) |
| KPCF | -0.36** | -0.33** | -0.37** | -0.43*** | -0.35** | -0.33** | -0.35** |
| (p-value) | (0.03) | (0.04) | (0.02) | (0.01) | (0.03) | (0.04) | (0.03) |

This table shows the Spearman correlations between GDP per capita and KACF2(KPCF). KACF2 means kurtosis of autocorrelation function with time lag 2. KPCF means kurtosis of the pair correlation function. ***, **, and * represent significant levels at 1%, 5%, and 10%, respectively. Values in parentheses are t-statistics. GDP represents the country's GDP per capita, which represents an indicator of a country’s standard of living.
